# Ethosuximide: subunit- and Gβγ-dependent blocker and reporter of allosteric changes in GIRK channels

**DOI:** 10.1101/2024.06.04.597296

**Authors:** Boris Shalomov, Theres Friesacher, Daniel Yakubovich, J. Carlo Combista, Haritha P. Reddy, Shoham Dabbah, Harald Bernsteiner, Eva-Maria Zangerl-Plessl, Anna Stary-Weinzinger, Nathan Dascal

## Abstract

The antiepileptic drug ethosuximide (ETX) suppresses epileptiform activity in a mouse model of *GNB1* syndrome, caused by mutations in Gβ_1_ protein, likely through the inhibition of G-protein gated K^+^ (GIRK) channels. Here we show that ETX is a subunit-selective, allosteric blocker of GIRKs. The potency of ETX block is increased by the G protein subunit dimer Gβγ, the physiological activator of GIRKs. Molecular dynamics (MD) simulations and mutagenesis locate the ETX binding site in GIRK2 to a region associated with phosphatidylinositol-4,5-bisphosphate (PIP_2_) regulation, and suggest that ETX acts by closing the HBC gate and altering channel’s interaction with PIP_2_. The apparent affinity of ETX block is highly sensitive to changes in channel gating caused by mutations in Gβ_1_ or GIRK subunits. Our findings pose GIRK as a potential therapeutic target for ETX, and ETX as a potent allosteric GIRK blocker and a tool for probing gating-related conformational changes in GIRK.

## Introduction

G-protein inwardly rectifying K^+^ channels (GIRK or K_ir_3) are important mediators of inhibitory neurotransmission in brain and heart, and are implicated in drug and alcohol addiction and several neurological disorders (Jeremic et al., 2021; Luo et al., 2022; Luscher & Slesinger, 2010). Neurotransmitters activate GIRKs via G-protein coupled receptors (GPCRs) by releasing the GIRK’s primary gating factor, Gβγ, from Gα_i/o_ (Dascal & Kahanovitch, 2015; Logothetis et al., 1987). A GIRK channel is a tetramer with four Gβγ-binding sites in its cytosolic domain (CSD), and two gates: the helix bundle crossing (HBC) formed by the transmembrane segments, and a cytosolic G-loop gate (Inanobe et al., 2007; Jin et al., 2002; Nishida & MacKinnon, 2002; Pegan et al., 2005) (Fig. 1A). Binding of Gβγ allosterically controls the gates, and potentially also the passage of ions through the selectivity filter (SF), in concert with the essential gating cofactor phosphatidylinositol 4,5-bisphosphate (PIP_2_), and sodium (Friesacher et al., 2022; Li et al., 2020; Sui et al., 1996; Sui et al., 1998; Wang et al., 2016). Maximal activation is attained when all four Gβγ-binding sites are occupied (Ivanova-Nikolova et al., 1998; Sadja et al., 2002; Wang et al., 2016; Yakubovich et al., 2015).

**Figure 1.**
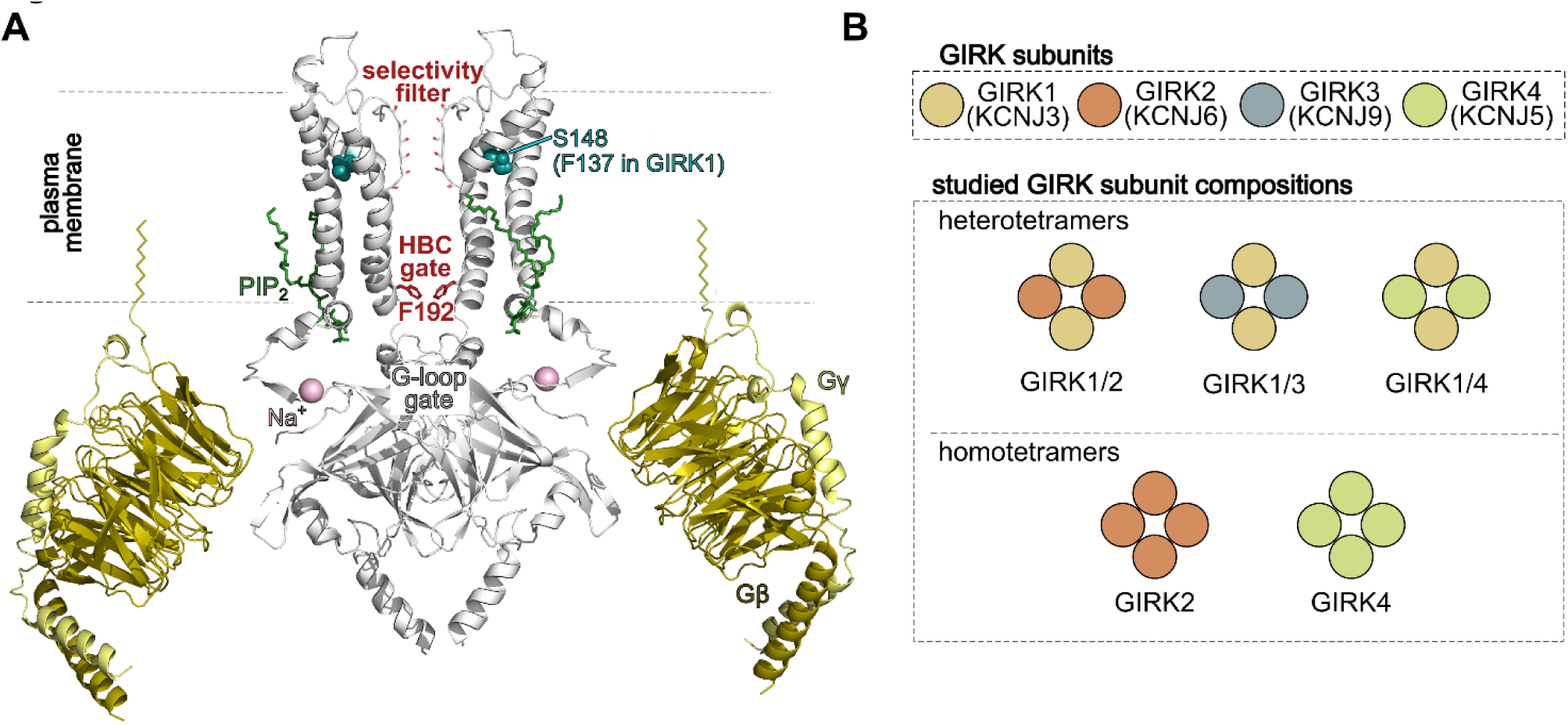
Structure of GIRK2 channel. **A**, side view of the GIRK2-Gβγ bound crystal structure (4KFM) (Whorton & MacKinnon, 2013). Two of the four GIRK2 subunits are shown in grey. The selectivity filter and the residue F192 composing the helix-bundle crossing (HBC) gate are colored in shades of red. The GIRK regulators PIP_2_, Na^+^ and Gβγ are shown in green, purple and yellow, respectively. The geranylgeranyl moiety (not in crystal structure), embedded into the plasma membrane, is shown schematically as yellow serpentine-like tail attached to the C-terminal end of Gγ. Residue S148 is presented in cyan spheres. **B**, schematic overview of the GIRK tetramers studied in this work. Gene names are shown below the GIRK acronyms used here. In the upper panel, gene names are shown along with standard subunit names used in the paper.

The four mammalian GIRK subunits form homotetrameric (GIRK2, GIRK4) or heterotetrameric (GIRK1/2, GIRK1/4, GIRK1/3, and GIRK2/3) channels. They vary in localization, gating properties, and plasma membrane (PM) expression (Jelacic et al., 2000; Ma et al., 2002; Rubinstein et al., 2009; Touhara et al., 2016). All are found in the brain, with GIRK1/2 being the most abundant (Lujan & Aguado, 2015). GIRK2 is the best-studied; several structures were published, but the exact allosteric pathway through which Gβγ activates GIRK channels remains uncertain (Mathiharan et al., 2021; Niu et al., 2020; Whorton & MacKinnon, 2013). GIRK1 has several distinctive features, notably a unique distal C-terminus (dCT) involved in the recruitment of Gβγ to the PM (Kahanovitch et al., 2014), and phenylalanine (F137) in the pore region, instead of a conserved serine in other GIRK subunits (Fig. 1A). Replacing F137 with serine enables the formation of functional homotetrameric GIRK1_F137S_ channels (termed GIRK1*) (Chan et al., 1996; Kofuji et al., 1996).

Due to their pivotal physiological role and their implication in disease, GIRK channels are considered promising targets for drug development (Jeremic et al., 2021). *GNB1* Encephalopathy is a neurodevelopmental disorder characterized by developmental delay and epilepsy, which is caused by mutations in the Gβ_1_ subunit (Lohmann et al., 2017; Petrovski et al., 2016; Revah-Politi et al., 2020). We found that certain disease-causing *GNB1* mutations alter Gβγ activation of GIRKs (Reddy et al., 2021). In a mouse model carrying the Gβ1 K78R mutation (which enhances Gβγ activation of GIRKs), epileptic symptoms were improved by ethosuximide (ETX), a known GIRK blocker (Kobayashi et al., 2009).

ETX is labeled as an absence seizure antiepileptic drug acting by blocking T-type voltage gated Ca^2+^ channels (VGCC) (Coulter et al., 1989; Gomora et al., 2001; Huguenard, 2002). However, therapeutic ETX doses (0.25-0.75 mM) do not strongly block T-type VGCCs, and the role of this mechanism in the antiepileptic action of ETX has been controversial (Crunelli & Leresche, 2002). Besides T-type VGCCs and GIRKs, ETX has been identified as an inhibitor of additional ion channels, including L-type VGCCs, non-inactivating Na^+^ currents, Ca^2+^-dependent K^+^ currents and IRK1 (Coulter et al., 1989; Crunelli & Leresche, 2002; Gomora et al., 2001; Huang & Kuo, 2015; Kobayashi et al., 2009).

In view of the potential role of GIRKs in the antiepileptic effect of ETX, we investigated the mechanism of ETX block of different GIRKs (Fig. 1B). Our findings reveal that ETX is a subunit-specific, potent, Gβγ-dependent allosteric blocker that inhibits neuronal Gβγ-activated GIRKs (GIRK1/2 and GIRK2) by 30-80% at therapeutic concentrations. Our findings unravel the molecular mechanisms of ETX action on GIRKs, and place GIRKs as potential primary ETX target, relevant to its antiepileptic action.

## Results

### ETX blocks GIRK channels in a subunit- and Gβγ-dependent manner and poorly blocks VGCCs

To characterize the effect of ETX on GIRK channels, we utilized the short isoform of mouse GIRK2 (mGIRK2), comprising 414 amino acids (a.a.), and the human GIRK2 (hGIRK2; 423 a.a.). We tested physiologically relevant subunit combinations in *Xenopus laevis* oocytes, with and without coexpression of Gβγ (Fig. 2; Fig. S1). GIRK currents and voltage-current (I-V) relationships were recorded in a high-K^+^ (24 mM) bath solution, HK24 (Rubinstein et al., 2009; Shalomov et al., 2022). ETX was applied in incremental concentrations, followed by Ba^2+^ to completely block GIRK currents (Fig. 2A,B). ETX dose-response relationships were fitted to Hill equation (Fig. 2C,D).

**Figure 2.**
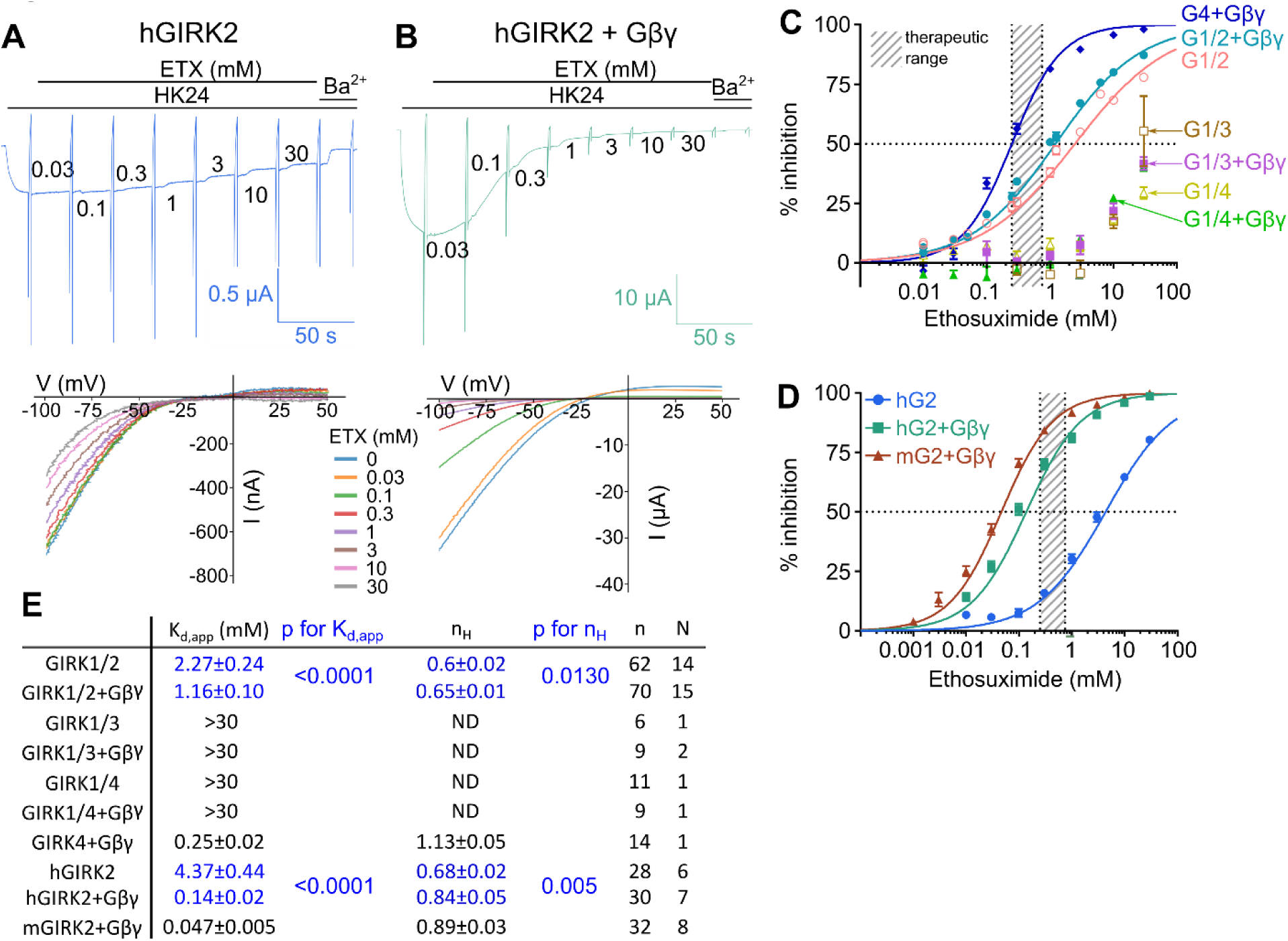
ETX block of GIRK channels is subunit- and Gβγ-dependent. Oocytes were injected with 5 ng Gβ_1_ RNA and 1 ng Gγ_2_ RNA. **A**, record of currents in a representative oocyte expressing hGIRK2 (top), and I-V relationships (bottom). GIRK currents were monitored using two-electrode voltage clamp. Holding potential was -80 mV. The physiological, 2 mM [K^+^]_out_ solution ND96, was switched to HK24 bath solution to observe inward GIRK currents. When the current reached steady-state, a voltage ramp from -120 to 50 mV was applied to obtain an I-V curve. Subsequently, the chamber was perfused with incremental concentrations of ETX (indicated, in mM, on the trace), with voltage ramps repeated for each concentration. Net GIRK currents were derived by subtracting currents remaining after blocking the GIRK channels with 2.5 mM Ba^2+^. **B**, same as **A** for an oocyte expressing hGIRK2+Gβγ. **C**, **D**, ETX dose response for different GIRK channel combinations across all experiments with all subunit combinations used (symbols present mean±SEM, number of experiments and cells is shown in Fig. 2E). G1 through G4 stand for GIRK1-GIRK4. For GIRK1/2, mGIRK2 was used. Fit of data to Hill equation was done in each cell, then K_d,app_ and n_H_ values were averaged across all cells and experiments. Solid lines for each subunit combination’s ETX dose-response were drawn using Hill equation with these average values. The striped rectangle shows the therapeutic range of ETX. **E**, mean±SEM of K_d,app_ and n_H_ obtained from Hill equation fits, from all experiments. For GIRK2 without Gβγ, ETX dose-response measurements were done only with hGIRK2, because (I_basal_) of mGIRK2 were too small (<100 nA). n = number of cells, N = number of experiments. Statistical analysis: effects of Gβγ on K_d,app_ and n_H_ of hGIRK2 and GIRK1/2 were compared by unpaired t-test for n_H_ of GIRK1/2 vs. GIRK1/2+Gβγ, Mann-Whitney test in all other cases.

In the absence of Gβγ, ETX inhibited hGIRK2 and GIRK1/2 channels with an apparent dissociation constant (K_d,app_) in the low-mM range (Fig. 2C-E). In contrast, ETX did not inhibit GIRK1/3 and GIRK1/4 at therapeutic doses. The sensitivity of GIRK1/3 to ETX was not studied so far; the other estimates align with prior research, except for GIRK1/4, which showed an IC_50_ of ∼1.5 mM (Kobayashi et al., 2009). This may reflect species-related differences; the rat GIRK4 used here and the mouse GIRK4 used by Kobayashi et al. (Kobayashi et al., 2009) differ by 13 a.a.

We have previously noticed that activation of GIRK1/2 and hGIRK2 by coexpression of Gβγ increased their apparent affinity (the reciprocal of K_d,app_) for ETX (Colombo et al., 2023). Here we compared the effect of channel activation by coexpressing a saturating concentration of Gβγ (Reddy et al., 2021; Yakubovich et al., 2015) (see Fig. 3A). The greatest shift in K_d,app_ was observed with GIRK2 (∼30 fold); more than 75% of the current was blocked by therapeutic doses of ETX (Fig. 2). Gβγ also modestly increased the apparent affinity for GIRK1/2 (∼2-fold), but did not affect the ETX block of GIRK1/3 or GIRK1/4 (Fig. 2C-E). The basal current (I_basal_) of homotetrameric GIRK4 channels was too small to measure K_d,app_, but Gβγ-activated GIRK4 showed high apparent affinity for ETX (Fig. 2C,E). The weak sensitivity of ETX block to Gβγ in GIRK1-containing heterotetramers, contrasting the high sensitivity of the homotetrameric GIRK2, suggests that the affinity of GIRK1 for ETX is not influenced by Gβγ. The Gβγ sensitivity of ETX block in GIRK1/2 likely arises from GIRK2.

**Figure 3.**
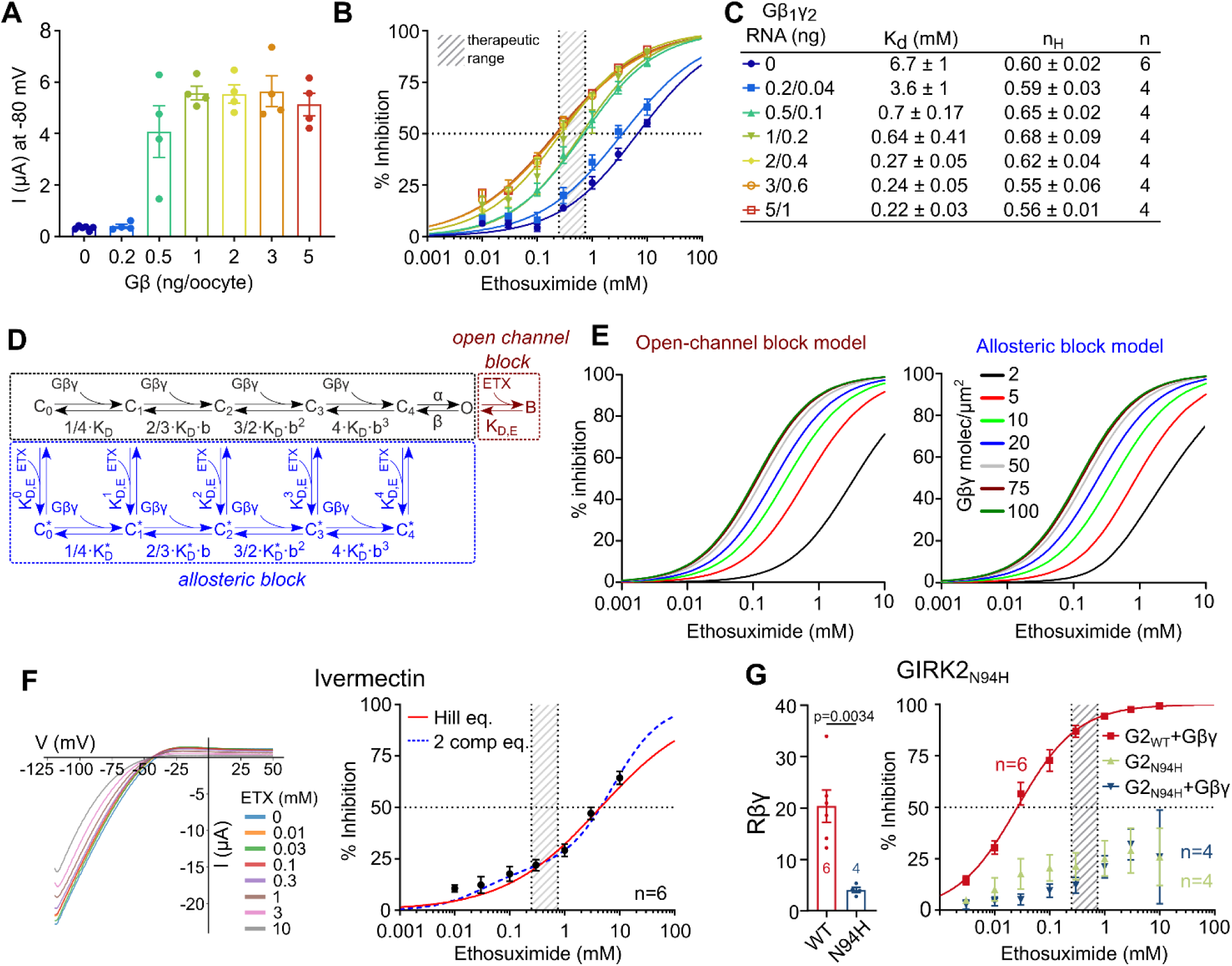
ETX block of GIRK2 is allosteric and Gβγ-dependent. **A-C** present the results of one experiment, representative of three. **A**, hGIRK2 (0.5 ng/oocyte) current amplitude at -80 mV, extracted from I-V curves, with increasing amounts of Gβγ. **B**, ETX apparent affinity is increased at higher expression levels of Gβγ. ETX dose-response relationships were tested in individual oocytes; symbols show mean % inhibition (±SEM), number of cells is shown in **C**, dose-response curves were fitted to Hill equation in each cell, and the average K_d,app_ and Hill coefficient (n_H_) values were used to draw the solid dose-response lines. **c** shows mean±SEM for K_d,app_ and n_H_, n is number of cells. **D**, two models of ETX block of GIRK2. Black: the basic scheme describing GIRK2 activation by Gβγ (Wang et al., 2016). Dark red: open channel block by ETX binding following channel opening. Blue: an allosteric model of ETX block. ETX can bind to closed states of the channel and its affinity is increasing with each additional Gβγ binding. α and β are the opening and closing rate constants for the transition C_4_ ⟺O. K_D_ and K_D,E_ are the dissociation constant of binding of Gβγ and ETX, respectively. K_D_* is the K_D_ of Gβγ binding to ETX-bound channel. Details on models are in Supplementary Methods, Fig. S3 and Table S1. **E**, simulated ETX dose-response curves of open channel block (left) and allosteric block (right) models. **F,** I-V curves of hGIRK2 activated by 2 µM IVM, at different doses of ETX, in a representative cell (left). Dose response of ETX block of hGIRK2 activated by 2 µM IVM is shown on the right. The dose response was fitted with Hill equation (Eqn. 2; K_d,app_=4.48±1.02 mM, n_H_=0.5±0.04) and a two component binding equation (Eqn. 3; K_1_=0.03±0.01 mM, K_2_=7.59±0.95 mM, c=0.21±0.03). **G**, comparison of Rβγ (fold activation by Gβγ) for GIRK2_WT_ and GIRK2_N94H_ reveals that the open channel N94H mutant was less sensitive to activation by Gβγ (left; unpaired t-test, p=0.0034). ETX dose response of GIRK2_N94H_ is shown on the right (K_d,app_=0.031±0.007 mM, n_H_=0.85±0.07).

We also assessed the impact of ETX on T-type VGCCs (Ca_V_3.2) and L-type VGCCs (Ca_V_1.2), highly expressed in the brain. Therapeutic doses of ETX only blocked less than 12% of the peak current of both channels (Fig. S2).

### ETX is an allosteric blocker of GIRK2

We next asked how the extent of Gβγ-induced activation of hGIRK2 affects the ETX block. Increasing the expression of Gβγ (by injecting incrementing amounts of Gβ and Gγ RNAs (Yakubovich et al., 2015)) enhanced GIRK2 currents (Fig. 3A) and, in parallel, reduced ETX’s K_d,app_ (Fig. 3B,C). Both processes saturated at about 1-2 ng Gβ_1_ RNA per oocyte, implying that fully Gβγ-activated channels have the highest ETX affinity.

The enhancement of ETX block upon GIRK2 activation by Gβγ indicates that ETX block is state-dependent. It could be a steric open-channel blocker that can enter the pore only after channel’s opening (Hille, 2002). Alternatively, the binding of ETX or its inhibitory action may be allosterically enhanced following Gβγ binding to GIRK2. To distinguish between these possibilities, we considered several models of ETX block (Fig. S3 and Table S1), based on the model developed by Wang et al. (Wang et al., 2016) (shown within the black rectangle in Fig. 3D). Here, Gβγ sequentially and cooperatively binds to the GIRK2 homotetramer; the channel opens when all four Gβγ are bound. Since the Hill coefficient (n_H_) for the ETX block of GIRK2 was ≤1, with or without Gβγ (Fig. 2e), we assumed a single binding site for ETX (Weiss, 1997). In state-dependent open or closed channel block models, ETX binds to the channel only in its open state (Fig. 3D, dark red square), or to a “preopen” state with four bound Gβγ (Fig. S3C). In a full allosteric scheme (Fig. 3D, blue square), ETX binds to each of the closed states occupied by 0 to 4 Gβγ molecules; the ETX binding affinity increases with each additional bound Gβγ. These models well simulated both the Gβγ-induced shift in ETX affinity and the range of K_d,app_ values for ETX block of GIRK2 without or with Gβγ (Fig. 3E, Fig. S3D).

In contrast, a model assuming ETX binding to any closed state with a constant, Gβγ-independent affinity did not predict the shift in K_d,app_ (Fig. S3E). Thus, ETX cannot act by indiscriminately binding to any channel’s state and must have a preference for a particular Gβγ-bound state(s). Altogether, although modeling did not distinguish between allosteric and steric block, it provided a framework for further investigation of the Gβγ-dependent increase in ETX apparent affinity.

We experimentally tested the open-channel block hypothesis using two approaches. First, we employed the Gβγ-independent activator of GIRK2 channels, ivermectin (IVM) (Chen et al., 2017). IVM concentration was adjusted to produce GIRK2 activation similar to Gβγ (Fig. S4A). Unlike Gβγ, IVM did not decrease K_d,app_ of ETX (Fig. 3F). In the second approach we used the constitutively open mutant mGIRK2_N94H_, which shows high activity in the absence of Gβγ (Yi et al., 2001). Compared to wild-type (WT) mGIRK2 (mGIRK2_WT_), mGIRK2_N94H_ had high I_basal_ (Fig. S4B) that was further increased by coexpressed Gβγ, but fold activation by Gβγ (Rβγ) was much less than for mGIRK2_WT_ (Fig. 3G). However, mGIRK2_N94H_ was very poorly blocked by ETX, with and without Gβγ (Fig. 3G). These results suggest that the apparent affinity of ETX block is not solely determined by channel’s opening (by any factor). Rather, it is channel’s activation by Gβγ that enhances the block, indicating that the effect of Gβγ is allosteric.

Interestingly, n_H_ smaller than 1 (0.6-0.7 in GIRK2 and GIRK1/2; Fig. 2E) indicates site heterogeneity (Colombo et al., 2023), implying two binding sites in a multimeric protein, or two populations of channels with different affinity (Ponstingl et al., 2005). The increase of n_H_ upon activation by Gβγ (Fig. 2E) indicates a higher fraction of high-affinity sites in the population, consistent with the allosteric model whereby the Gβγ-induced conformational change enhances ETX interaction with the channel.

### Gβγ effect on ETX affinity is absent in loss-of-function Gβ_1_ mutants

We have previously characterized several mutants of Gβ_1_ that cause the *GNB1* Encephalopathy (Colombo et al., 2023; Reddy et al., 2021). Gβ_(I80N)_ and Gβ_(I80T)_ showed loss-of-function (LoF) toward homotetrameric GIRK2 but activated GIRK1/2, probably through GIRK1. Gβ_(K78R)_ showed gain of expression and gain of function (GoF) toward GIRK2 and GIRK1/2, but impaired GIRK1/2 expression and gating when expressed at high levels (Colombo et al., 2023; Reddy et al., 2021). We tested how the three Gβ mutants affect the dose dependence of ETX block. All mutants and Gβγ_WT_ equally increased GIRK1/2 currents (Fig. 4A). Both Gβγ_WT_ and Gβ_K78R_γ comparably decreased K_d,app_ of ETX block of GIRK1/2, as reported previously (Colombo et al., 2023) (Fig. 4B,C). In contrast, despite the equivalent activation (opening) of the channel, neither Gβ_I80N_γ nor Gβ_I80T_γ enhanced the ETX block; Gβ_I80T_γ even reduced the apparent ETX affinity (Fig. 4B,C). These results support the allosteric nature of regulation of ETX block by Gβγ.

**Figure 4.**
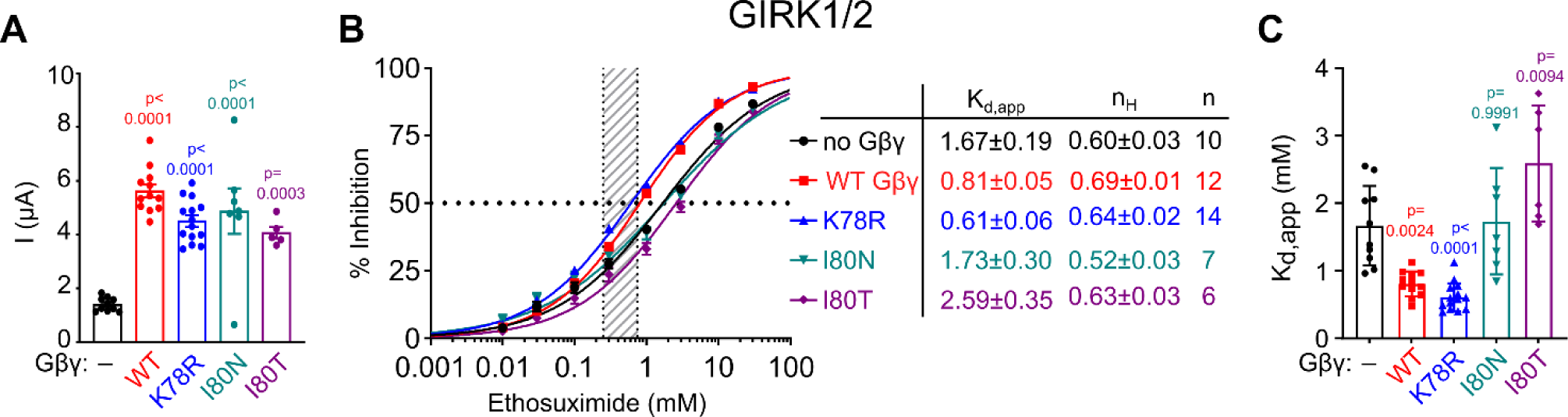
Loss-of-function mutants of Gβ lose the ability to regulate ETX blocking effect in GIRK1/2 channels. Results of one experiment are shown, representative of two. Oocytes were injected with 50 pg RNA of both GIRK1 and mGIRK2 subunits, and 5 ng Gβ RNA and 1 ng Gγ RNA (when present). **A**, GIRK1/2 channels are similarly activated by Gβγ_WT_ and the Gβ_1_ mutants K78R, I80N, and I80T. The p values above bars relate to the difference from control (I_basal_, no Gβγ). There was no difference among Gβγ-activated groups (p>0.05 in all pairwise comparisons; one-way ANOVA followed by Tukey’s test). **B**, ETX dose response of GIRK1/2 without or with WT or mutant Gβγ (left), and the Hill plot parameters from fits in individual oocytes (right). Mean±SEM values are listed. **C**, comparison of K_d,app_ values extracted from data shown in **B** show that Gβγ_WT_ and Gβ_K78R_γ increased the apparent affinity of ETX, whereas Gβ_I80N_γ did not change, and Gβ_I80T_γ decreased the apparent ETX affinity (one-way ANOVA followed by Tukey test). The values of p vs. control (no Gβγ) are shown.

### The binding site of ETX to GIRK2 channel lies in the transmembrane domain

Since the apparent affinity of ETX block of GIRK1/3 was very low and Gβγ-insensitive (Fig. 2C, E), it seemed plausible that GIRK3 does not bind ETX. As a first step in locating the ETX binding site, we constructed GIRK2-GIRK3 chimeras with swapped transmembrane (TMD) and cytosolic (CSD) domains (GIRK2_G3TM_ and GIRK3_G2TM_; Fig. 5A). Notably, the TMDs of GIRK2 and GIRK3 differ by 24 a.a. (Fig. 5B). The chimeras did not produce measurable currents when expressed as homotetramers. Therefore, we coexpressed them with GIRK1, which produced functional channels, albeit with a reduced activity compared to GIRK1/2_WT_ (Table S2; note the different RNA amounts used). Nonetheless, Gβγ activation was comparable or better than for GIRK1/2_WT_ (Fig. 5C). Importantly, we observed that GIRK1/2_G3TM_ lost sensitivity to ETX, whereas GIRK1/3_G2TM_ gained sensitivity to ETX (Fig. 5C,D). These results indicate that ETX binds to the TMD of GIRK2.

**Figure 5.**
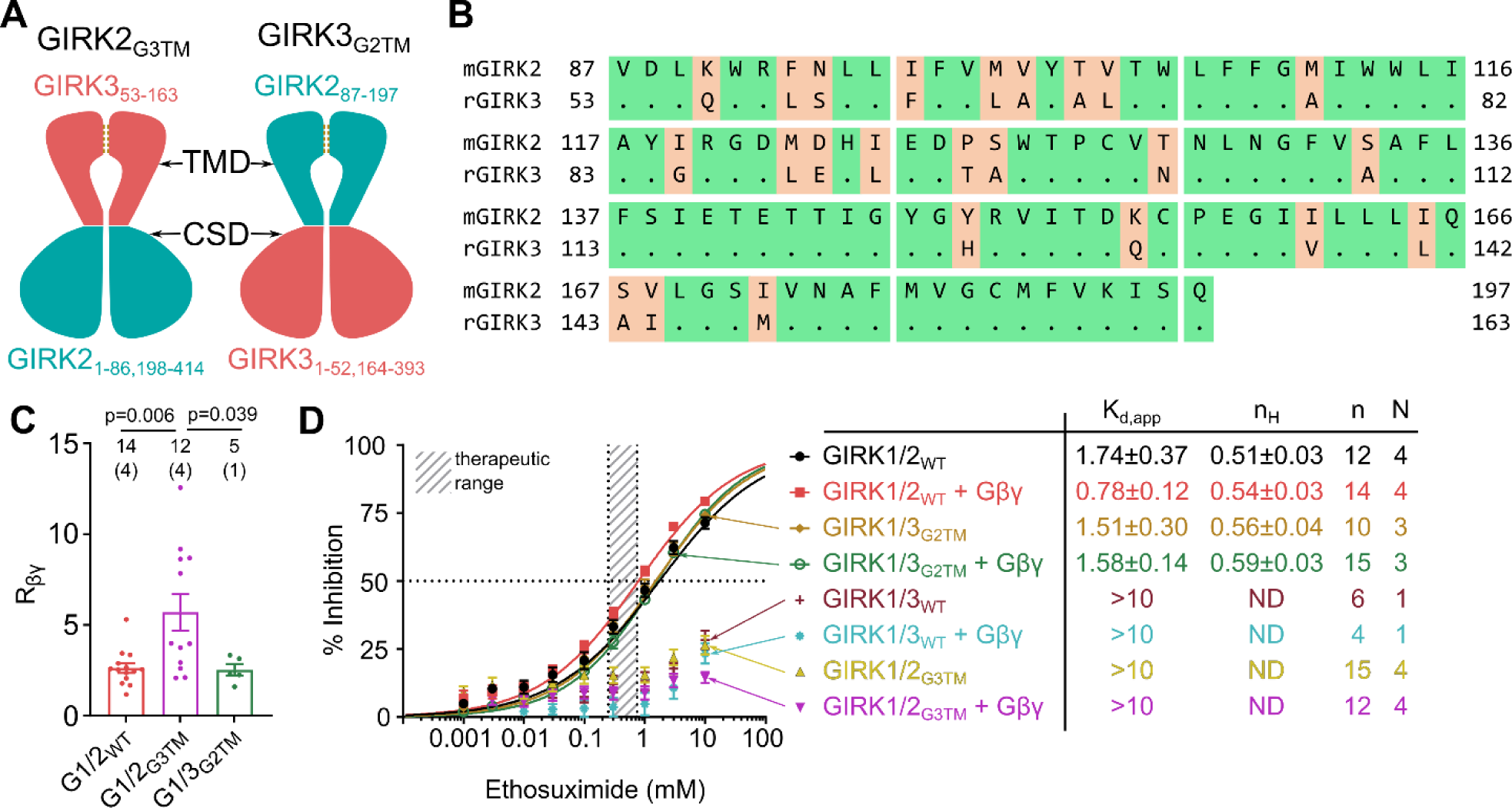
GIRK3 gives clues for the ETX binding site. **A**, the GIRK2-GIRK3 chimeras used. Note that in GIRKs the CSDs consist of a short N-terminal domain and a long C-terminal domain. **B**, sequence alignment of TM region from GIRK2 and GIRK3. Dots indicate identical a.a. **C**, comparison of the extent of Gβγ activation. One-way ANOVA (F (3, 46) = 75.17, p<0.0001) followed by Tukey test. **D**, ETX dose response results. GIRK1/2_G3TM_ lost the sensitivity to ETX and became similar to GIRK1/3, while GIRK1/3_G2TM_ gained sensitivity to ETX, becoming similar to GIRK1/2_WT_. Details on RNA doses used, current amplitudes and Rβγ found in these experiments are shown in Table S2.

### MD simulations reveal a putative ETX binding site in GIRK2 and an ETX-induced narrowing of the HBC gate

To investigate GIRK inhibition by ETX, we carried out MD simulations with the open and conductive GIRK2 channel bound to PIP_2_ (PDB:3SYA) (Whorton & MacKinnon, 2011) from our previous study (Bernsteiner et al., 2019). Twenty ETX molecules were placed randomly in the solvent, resulting in a concentration of 18.2 mM. We conducted 5 replicas (run 1-5) of 1.5 μs-long unbiased MD simulations. In run1, run2 and run3, we observed that ETX binds to one of the GIRK subunits close to the activator PIP_2_ (Fig. 6A,C) for the duration of 1.2 μs, 0.7 μs and 0.6 μs, respectively (Fig. 6B).

**Figure 6.**
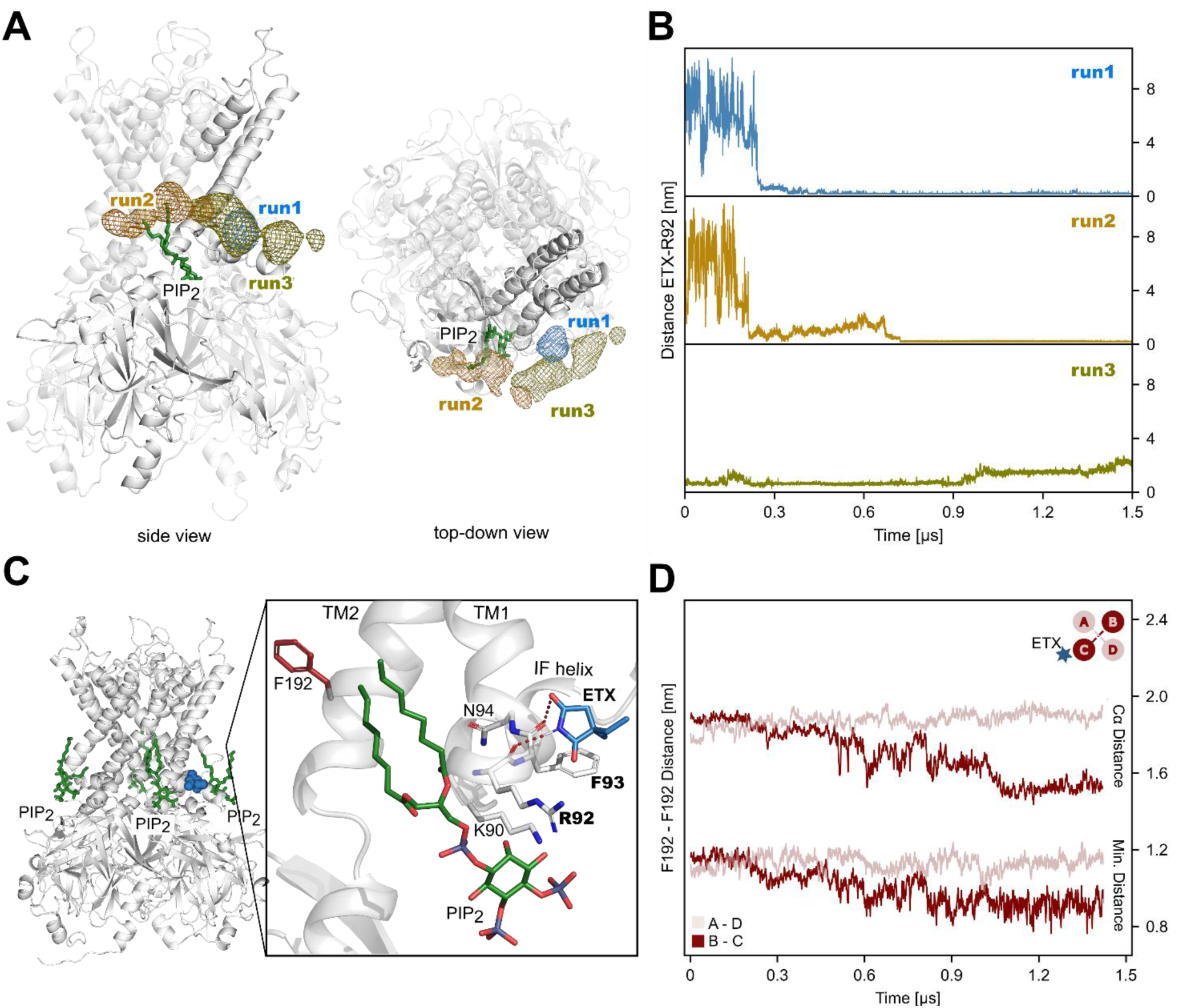
ETX binds to the GIRK2 channel close to the activator PIP_2_ and leads to closing of the HBC gate. **A**, aligned time-averaged density maps of ETX molecules binding to GIRK2 near the activator PIP_2_. The maps are shown as blue, orange, and olive mesh for run1, run2, and run3, respectively. PIP_2_ is colored green, and one subunit of GIRK2 is highlighted in grey. **B,** distances between the ETX molecules binding to GIRK2 near the activator PIP_2_ and the GIRK2 residue R92. **C,** ETX binding site identified in run1 during 1.5 μs free MD simulations. On the right, a close-up of the ETX binding site is displayed. The GIRK2 channel, PIP_2_, and ETX are colored in grey, green, and blue, respectively. **D,** minimum distances and Cα distances between the HBC gate-forming F192 of opposing subunits in run1. Pairs of opposing subunits are colored in different shades of red. ETX binds to GIRK subunit C.

A closer look reveals that the ETX-binding region is located around residue R92 (Fig. 6A,B). ETX adopts multiple binding poses, before settling into a stable conformation (Fig. 6B,C; in run1 after 370 ns). Remarkably, we observe a concomitant narrowing of the HBC gate (Fig. 6D). The minimum distance between the opposing F192, which compose the constriction point of the HBC gate, decreases by 2 Å, while the distance between the Cα atoms drops by 5 Å. Hydrogen bonds are formed between the amino and carbonyl moieties of the drug and the backbone oxygens of R92 and F93 of GIRK2 (Fig. 6C). ETX is stacked against the TM1 of one subunit at the level of the HBC gate within 10 Å of PIP_2_.

### The proposed allosteric mechanism of GIRK2 inhibition by ETX

A thorough investigation of the channel dynamics revealed that, in run1, the ETX-bound GIRK2 subunit experiences an outward swing and rotation of the outer helix TM1, accompanied by an inward movement and rotation of pore-lining helix TM2 that harbors the HBC gate residue F192 (Fig. S5). To investigate how the conformational change in the TM1 affects the TM2, we extracted shortest pathways between the ETX-binding residues R92 and F93, and the HBC gate residue F192 with the “get Residue Interaction eNergies and Networks” (gRINN) tool (Sercinoglu & Ozbek, 2018). We revealed a chain of interacting amino acids R92-W91-K194-F192 (Fig. 7A). Upon ETX binding, the intracellular end of the TM1 (K90-F93) is displaced, changing the side chain torsion angle χ_1_ of W91 (Fig. 7B). The new side chain orientation enables a better interaction between W91 and the TM2 residue K194, which is pushed towards the helix pore, thereby displacing F192, and closing the HBC gate (Fig. 7). Another pathway coupling the ETX binding site to F192 is via F93-L89-K194-F192 (Fig. 7A). F93 transfers the drug binding-induced conformational change to L89 (a part of the interfacial helix), leading to an enhanced interaction between L89 and K194 (Fig 7b). In addition to changes in the intra-residue interactions, ETX binding to GIRK is accompanied by a displacement of PIP_2_ from its binding site (Fig. S6).

**Figure 7.**
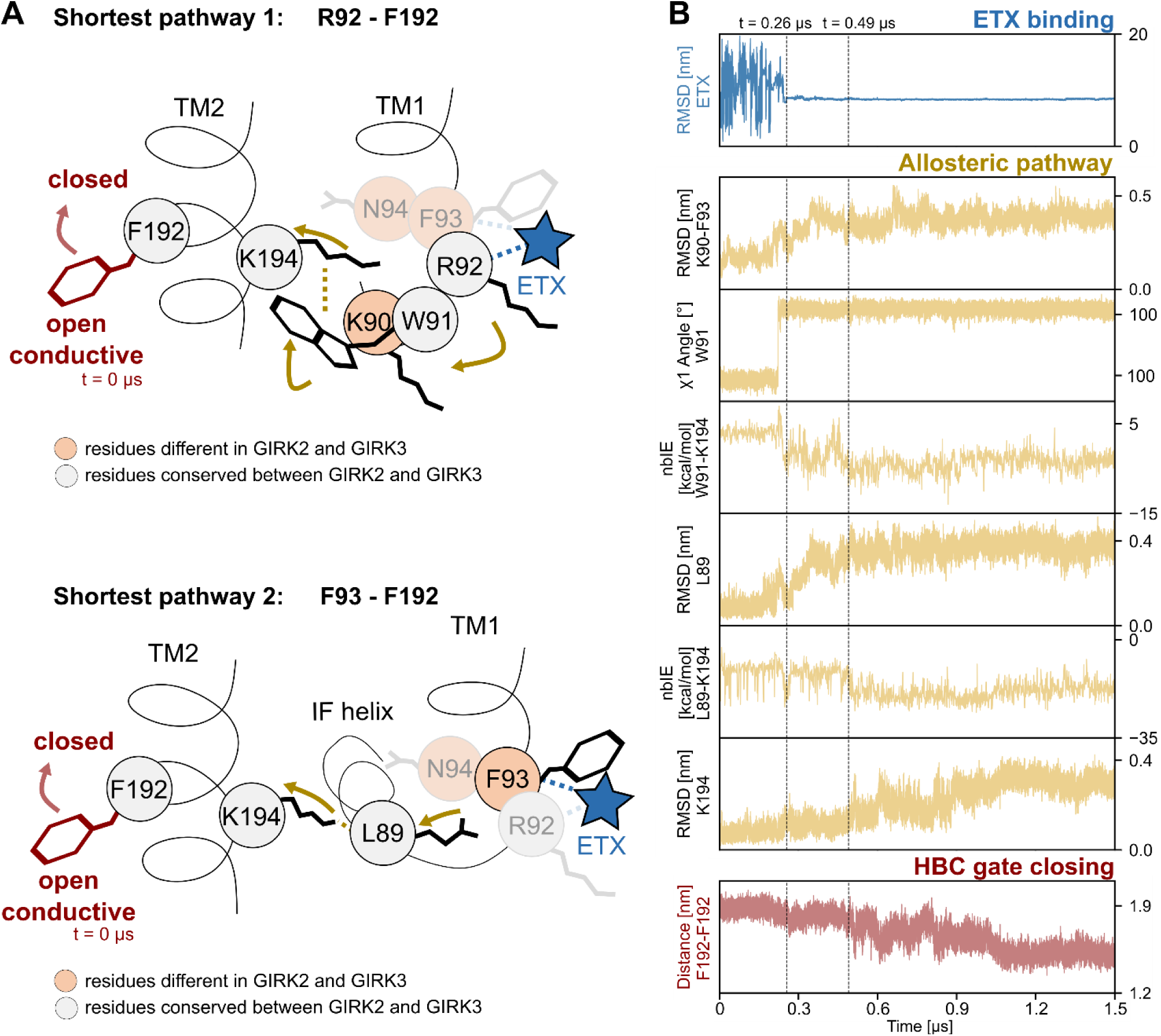
Shortest pathways between the ETX binding site and the HBC gate based on the protein network analysis. **A,** schematic illustration of the shortest pathways between the ETX binding residues R92 and F93, and the HBC gate residue F192. ETX and interactions of ETX with GIRK2 are shown in blue, while F192 and conformational changes of this residue are shown in red. Other GIRK2 residues are shown in grey, while conformational changes and interactions between these residues are shown in yellow. Arrows show conformational changes, while dotted lines indicate non-bonded interactions between residues. **B**, over-time plots showing stable interactions between ETX and GIRK in blue, the resulting conformational changes and alterations of non-bonded interactions energies (nbIE) of channel residues in yellow, and HBC gate closing in red.

The network betweenness centrality (BC) indicates how efficiently a residue passes on information, while the closeness centrality (CC) determines the importance of the residue for information transfer. Ranking GIRK2 residues by their BC and CC scores reveals that the PIP_2_ binding residue K199 and L89, located on the interfacial helix, are key players in the network. Furthermore, N94, which had the second highest BC score, is a leading residue for information transfer.

### Mutagenesis of amino acids around R92 reduces the affinity of ETX to GIRK2

MD simulations highlighted the significance of amino acid residues near GIRK2_R92_, specifically K90-N94. K90-N94 are conserved in GIRK4 but GIRK3 has three mismatches and GIRK1 has one mismatch (Fig. 8A). Notably, GIRK2_R92_ is central in channel’s interaction with PIP_2_ (Whorton & MacKinnon, 2011). Arginine at this position is conserved in all GIRK subunits (Fig. 8A) and nearly all Kir channels. Therefore, our approach involved retaining the conserved W91 and R92 while introducing reciprocal mutations in a.a. residues that distinguish GIRK2 from GIRK3. We produced GIRK2 with the substitutions K90Q, F93L, and N94S, and a triple mutant, GIRK2_QLS_. In GIRK3, the substitutions were reversed: Q56K, L59F, and S60N, and the triple mutant GIRK3_KFN_.

**Figure 8.**
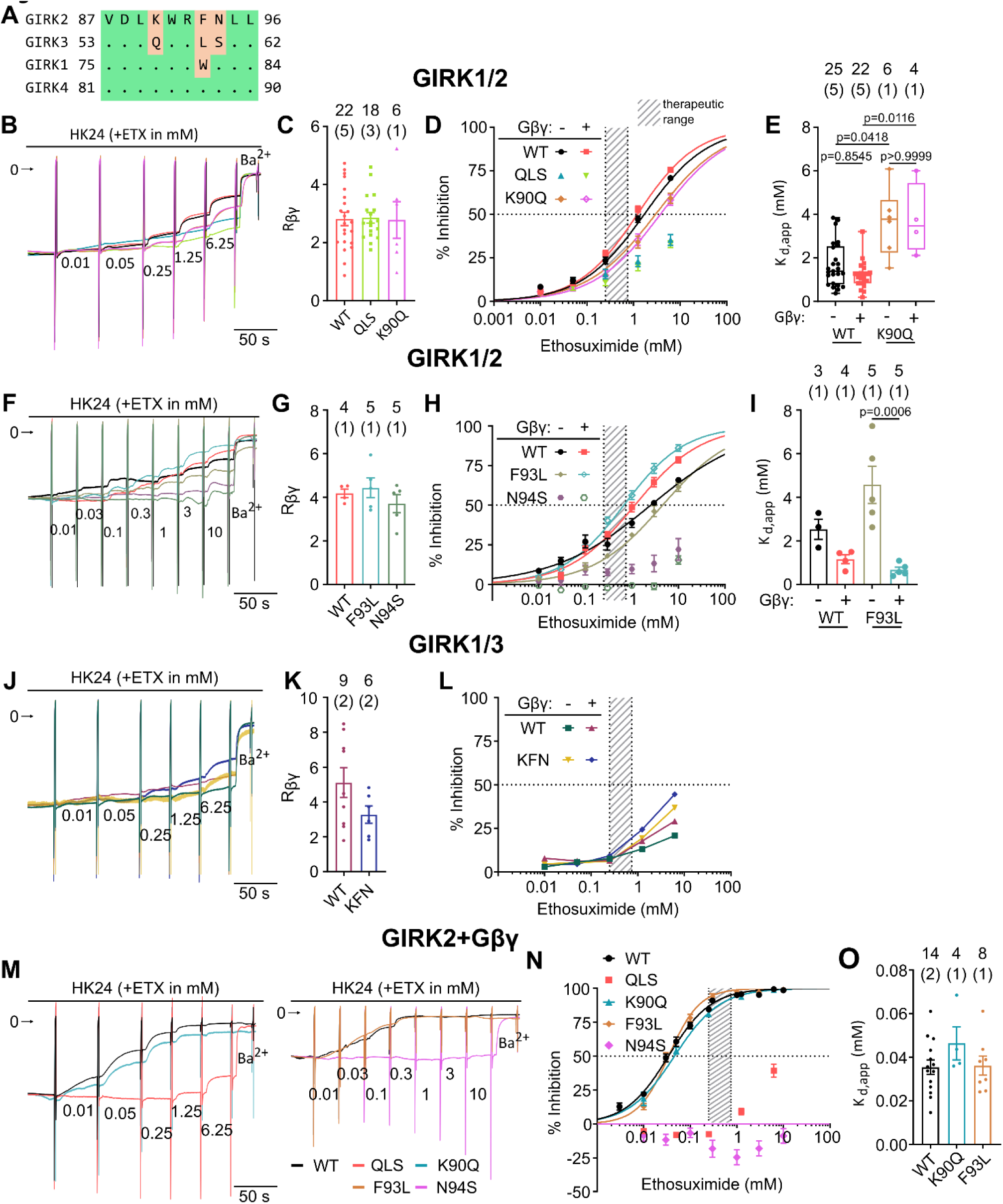
Mutagenesis in the PIP_2_ binding region reveals possible ETX binding site. **A**, sequence alignment of the PIP_2_ binding region revealed disparities between all GIRK subunits. **B**, ETX effect on currents of GIRK1/2_WT_, GIRK1/2_QLS_, and GIRK1/2_K90Q_ with or without Gβγ. Currents were normalized to their magnitudes before ETX addition. **C**, GIRK1/2_QLS_ and GIRK1/2_K90Q_ are activated by Gβγ like GIRK1/2_WT_. **D**, ETX dose response of GIRK1/2_WT_, GIRK1/2_QLS_, and GIRK1/2_K90Q_ with or without coexpressed Gβγ show that GIRK1/2_QLS_ becomes less sensitive to ETX. **E**, K_d,app_ extracted from 8D show that the GIRK1/2_K90Q_ was less sensitive than GIRK1/2_WT_ (Kruskal-Wallis test followed by Dunn’s multiple comparison test). **F**, normalized recordings of ETX effect on GIRK1/2_WT_, GIRK1/2_F93L_, and GIRK1/2_N94S_ with or without Gβγ. **G**, Rβγ of GIRK1/2_F93L_ and GIRK1/2_N94S_ is similar to GIRK1/2_WT_. **H**, ETX dose responses of GIRK1/2_WT_, GIRK1/2_F93L_, and GIRK1/2_N94S_ with or without coexpressed Gβγ show that GIRK1/2_N94S_ becomes less sensitive to ETX. **I**, K_d,app_ extracted from 8H (One-way ANOVA followed by Tukey’s test). **j**, normalized recordings of ETX effect on GIRK1/3_WT_ and GIRK1/3_KFN_ with or without Gβγ. **K**, Rβγ of GIRK1/3_KFN_ is similar to GIRK1/3_WT_. **L**, ETX dose response of GIRK1/3_WT_ and GIRK1/3_KFN_ with or without Gβγ. **M**, normalized recordings of ETX effect on GIRK2_WT_, GIRK2_QLS_, GIRK2_K90Q_, GIRK2_F93L_ and GIRK2_N94S_ coexpressed with Gβγ. Left and right panels show two separate experiments. **N**, ETX dose responses of homotetrameric GIRK2_WT_ and mutants (from 2 experiments shown in m). **O**, K_d,app_, summary of the two experiments. Full details on Hill equation fit parameters for all experiments are shown in Table S3.

We started with GIRK1/2 and GIRK1/3 heterotetramers (since GIRK3 homotetramers are not functional), and employed the GIRK3_Y350A_ mutant lacking the lysosomal-targeting motif (Ma et al., 2002) to improve PM expression. First, we compared GIRK1/2_QLS_ and GIRK1/2_K90Q_ to GIRK1/2_WT_ (Fig. 8B-E). Gβγ activated GIRK1/2_QLS_ and GIRK1/2_K90Q_ like GIRK1/2_WT_ (Fig. 8C, Fig. S7A,B). However, GIRK1/2_QLS_ showed a significant reduction in sensitivity to ETX, even when coexpressed with Gβγ (K_d,app_ > 10 mM). For GIRK1/2_K90Q_, the ETX affinity was 2-fold lower without Gβγ and 3-fold lower with Gβγ, compared with GIRK1/2 (Fig. 8D,E). The mutants GIRK1/2_F93L_ and GIRK1/2_N94S_ were also activated by Gβγ and showed Rβγ comparable to GIRK1/2_WT_ (Fig. 8g; Fig. S7C,D). The K_d,app_ of ETX block in GIRK1/2_F93L_ was similar to GIRK1/2_WT_, and coexpression of Gβγ significantly reduced K_d,app_. In contrast, GIRK1/2_N94S_ was insensitive to ETX, irrespective of Gβγ expression (Fig. 8H,I).

We next tested the reciprocal triple mutation in GIRK3 (Fig. 8J-L, Fig. S7E,F). GIRK1/3_KFN_ was activated by Gβγ similarly to GIRK1/3_WT_. The block by ETX appeared stronger than in GIRK1/3_WT_, though still less than 50% at 10 mM ETX. Altogether, the exchange between GIRK2 and GIRK3 of the three a.a. in the immediate vicinity of R92 (GIRK2 count), in the GIRK1/x heterotetrameric context, causes corresponding shifts in the apparent ETX affinity.

We further studied the effect of the QLS mutations in the context of homotetrameric mGIRK2 channels. Because of mGIRK2’s low I_basal_, Gβγ was coexpressed in all cases. Gβγ-activated mGIRK2_QLS_ and mGIRK2_K90Q_ had significantly lower activity than mGIRK2_WT_ (Fig. S7G,H). However, mGIRK2_K90Q_ had a similar apparent affinity for ETX as mGIRK2_WT_, whereas mGIRK2_QLS_ showed a significant loss of affinity to ETX, with no block up to 1 mM ETX (Fig. 8M-O). For the individual F93L and N94S mutations, the Gβγ-evoked currents were similar to mGIRK2_WT_ (Fig. S7I,J). mGIRK2_F93L_ showed a similar affinity for ETX as mGIRK2_WT_. Remarkably, mGIRK2_N94S_ was completely insensitive for ETX (Fig. 8M-O), as was seen also with heterotetrameric with GIRK1/2_QLS_ and GIRK1/2_N94S_ and with the previously tested mutant of N94, N94H (Fig. 3G). These results support the predictions of the MD analysis, suggesting the importance of a.a. 90-94 of GIRK2 for ETX binding and (in particular N94) for its allosteric link to channel gating.

### The role of S148 of GIRK2 and F137 of GIRK1 in the allosteric effect of ETX

The a.a. residues crucial for ETX effect in GIRK2 are conserved in GIRK1 except GIRK2_F93_, which is a tryptophan (W82) in GIRK1. However, a similar homologous substitution, F93L, did not affect ETX block in GIRK2 (Fig. 8). According to this logic, GIRK1 should bind ETX. The low sensitivity of ETX block to Gβγ in GIRK1/2 channel could result from a deficiency in allosteric coupling between ETX block and Gβγ in GIRK1.

To better understand if GIRK1 binds ETX, we utilized the GIRK1* homotetramers formed by the GIRK1_F137S_ mutant, GIRK1* (Chan et al., 1996; Kofuji et al., 1996). GIRK1* is activated by Gβγ but shows single channel conductance (i_single_) and open probability (P_o_) of about 50% of those of GIRK1/4 (Chan et al., 1996) or GIRK1/2 (Fig. S8). We also noticed that GIRK1* is less selective to K^+^ than GIRK1/2, showing a ∼15 mV depolarized shift in the reversal potential (V_rev_) (Fig. S9). These observations suggest that the open pore conformation of GIRK1* significantly differs from GIRK1/2 or GIRK1/4.

Strikingly, ETX blocked the GIRK1* channel with a remarkably high apparent affinity, comparable to Gβγ-activated GIRK2 (Fig. 9A). Coexpression of Gβγ increased the current and further increased the affinity for ETX ∼3 fold (Fig. 9A, Fig. S8B). Clearly, GIRK1* binds ETX. The possibility that serine at this position itself confers ETX binding (and is part of an ETX-binding site) is highly unlikely, because i) this serine is present in the ETX-insensitive GIRK3; ii) in GIRK2, S148 is not solvent-accessible from the external solution or from within the membrane (Fig. 10A). We conclude that GIRK1_WT_ also contains an ETX binding site. We propose that serine at position 137 in GIRK1 (analogous to S148 in GIRK2) enables the allosteric pathway, present in GIRK2 and GIRK4, that links between ETX binding, channel block, and Gβγ activation. In line with this, network analysis in GIRK2 identifies a pathway linking S148 with N94 (ETX sensitive), and R337 (part of the Gβ-binding site (Whorton & MacKinnon, 2013)) (Fig. 10B).

**Figure 9.**
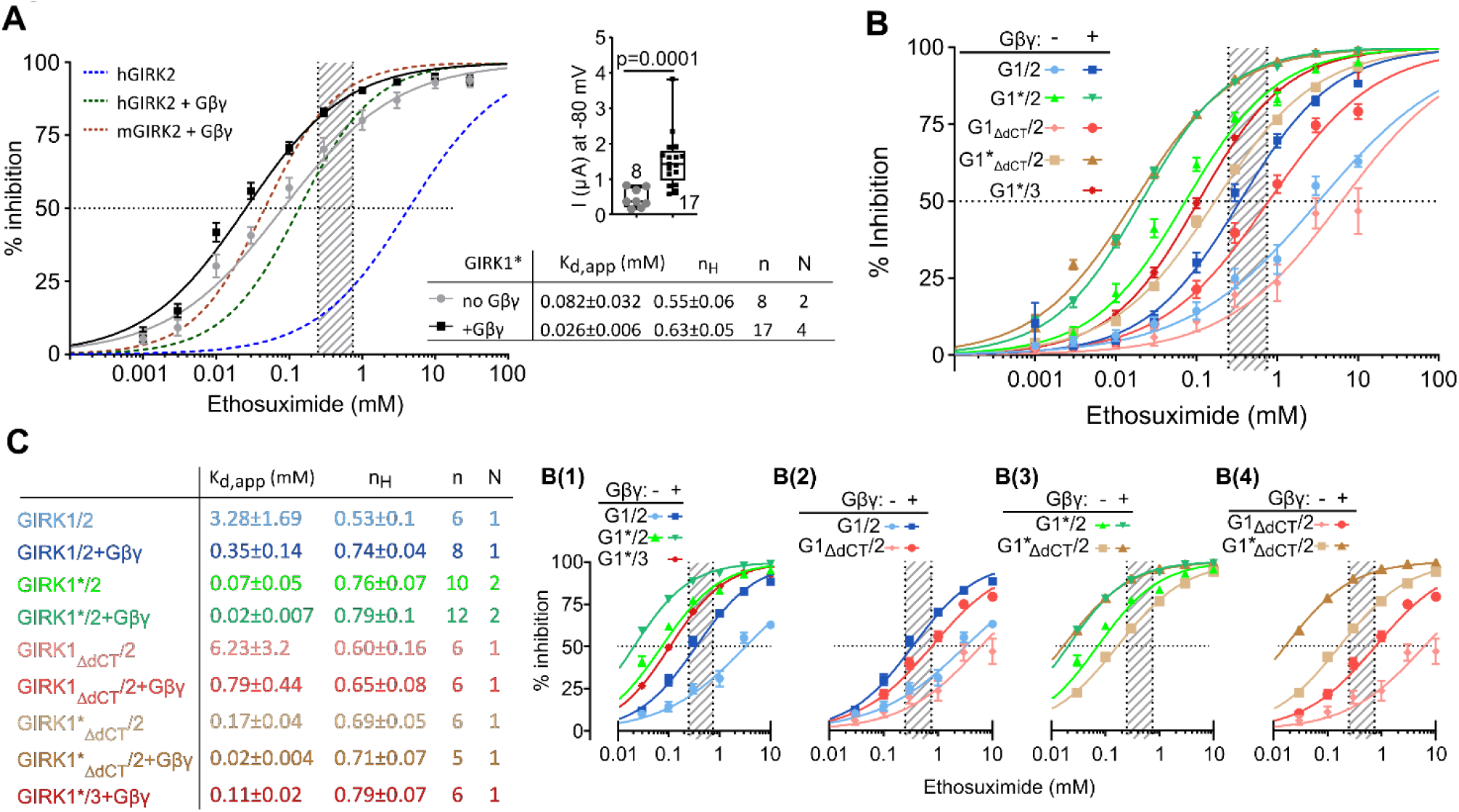
ETX uncovers the role of F137 of GIRK1 in the allosteric pathway of channel gating. **A,** ETX strongly blocks GIRK1*, with or without coexpressed Gβγ. The dotted lines show the ETX dose-response curves of hGIRK2, hGIRK2+Gβγ, and mGIRK2 from Fig. 2D. The striped rectangle shows the therapeutic range of ETX. **B**, ETX dose-response of GIRK1/2, GIRK1*/2, GIRK1_ΔdCT_/2, and GIRK1*_ΔdCT_/2 with and without coexpressed Gβγ, and GIRK1*/3 with coexpressed Gβγ. Panels **B(1)-B(4)** zoom on subsets of dose-response curves. Notation: G1/2 stands for GIRK1/2, G1*/2 stands for GIRK1*/2, and so on. **C**, K_d,app_ and n_H_ extracted from 9B.

**Figure 10.**
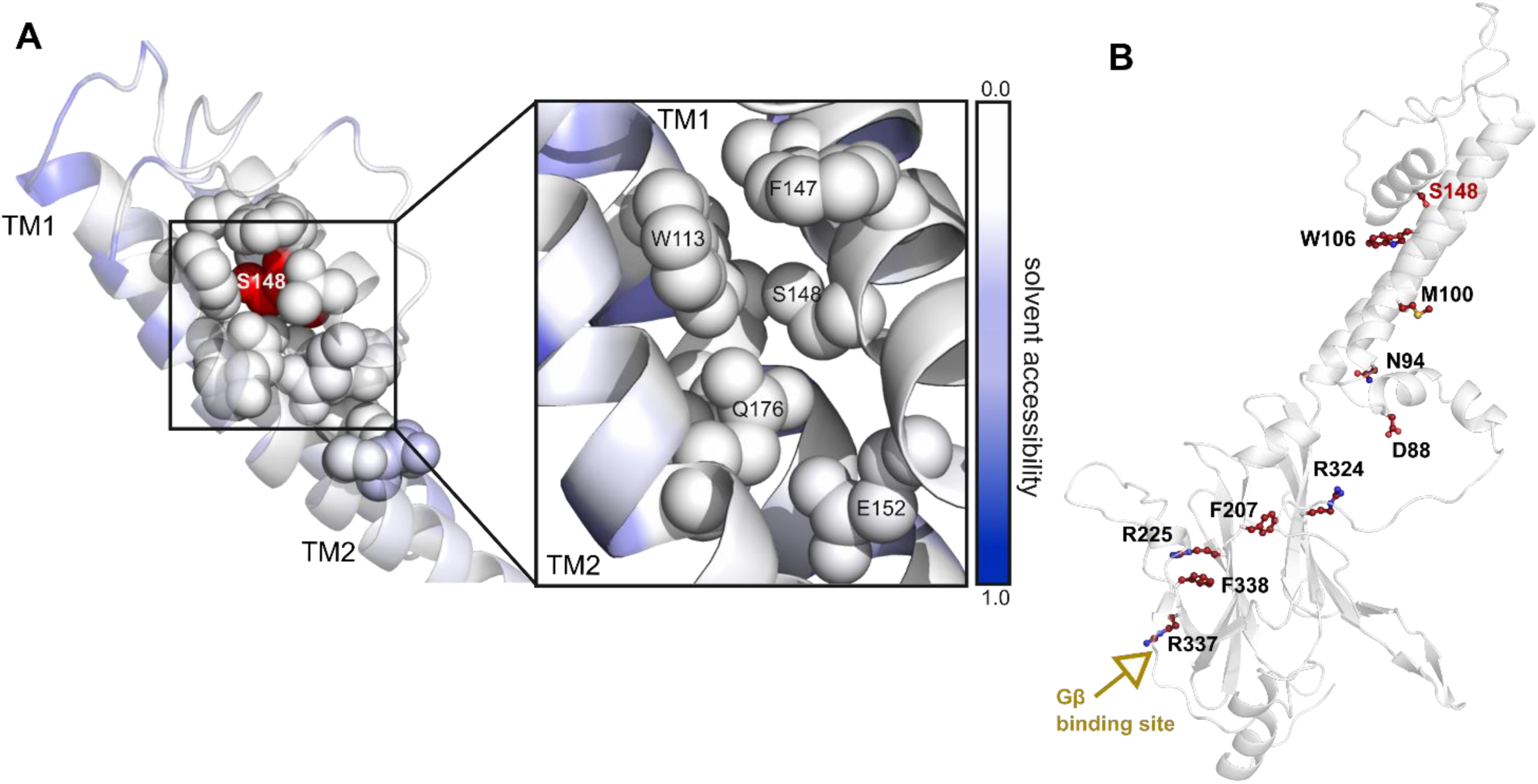
S148 is not solvent-accessible but is part of the allosteric network coupling ETX- and Gβγ-binding sites, along with N94, in GIRK2. **A**, S148 in GIRK2 structure. S148 and the surrounding residues are represented as spheres. The left figure shows the upper part of the TMD, with S148 colored red. On the right, a close up of S148 and its surrounding residues can be seen. The structures are colored according to their local solvent accessibility, with white and purple meaning no and full solvent accessibility, respectively. **B**, the shortest pathway linking S148 and the Gβ binding residue R337 passes through the GIRK2 residue N94, important for channel inhibition by ETX. GIRK2 is shown as white cartoon, the residues of the pathway are represented as red sticks. The network was calculated based on the dynamics of subunit C in run1 that binds ETX in the course of the MD simulation.

GIRK1* and GIRK1/2 recruit Gβγ owing to the presence of a high-affinity Gβγ-binding “anchoring” site in GIRK1 (Dascal & Kahanovitch, 2015). GIRK1’s unique dCT (GIRK1-dCT) is an essential component of this site, and its removal abolishes the Gβγ recruitment and reduces the I_basal_ of GIRK1* and GIRK1/2 in *Xenopus* oocytes (Kahanovitch et al., 2014; Rubinstein et al., 2009). Could the high ETX affinity of GIRK1* be due to channel’s activation by pre-associated Gβγ? To address this possibility, we employed GIRK1 and GIRK1* lacking the last 121 a.a., GIRK1_ΔdCT_ and GIRK1*_ΔdCT_ (Kahanovitch et al., 2014). As homotetrameric GIRK1*_ΔdCT_ gives very small currents (Tabak et al., 2019), we focused on heterotetrameric combinations with mGIRK2_WT_ and GIRK3.

All combinations of GIRK1_ΔdCT_ and GIRK1*_ΔdCT_ constructs with GIRK2 had measurable I_basal_, were activated by Gβγ (Fig. S10), and Gβγ caused a pronounced shift in K_d,app_ of ETX, suggesting an intact regulation of ETX affinity by Gβγ (Fig. 9B,C). The removal of GIRK1-dCT resulted in a ∼2-2.5-fold reduction in apparent ETX affinity both in GIRK1/2 and in GIRK1*/2 (increase in K_d,app_ from 3.3 to 6.2 mM and from 0.07 to 0.17 mM, respectively; Fig. 9B2,C). After coexpression of Gβγ, K_d,app_ was identical in GIRK1*/2 and GIRK1*_ΔdCT_/2 (∼0.02 mM; Fig. 9B3,c). The K_d,app_ of GIRK1_ΔdCT_/2 (0.79±0.44 mM, n=6, Fig. 9B2,C) was also similar to the average K_d,app_ for GIRK1/2 across all experiments (1.22±0.09, n=82, p=0.166, compare Fig. 9C and Fig. 2E). These results demonstrate that GIRK1-dCT *per se* does not regulate the ETX block, and rule out the possibility that the high apparent affinity of GIRK1* to ETX results from a pre-docked Gβγ. Rather, the pre-associated endogenous Gβγ appears to mildly increase ETX apparent affinity in GIRK1-containing channels in basal state, but not after saturation by Gβγ.

It seemed logical to explore the swapped mutant, GIRK2_S148F_, to further address the role of F148 of GIRK2. The results were inconclusive. Heterotetramers of GIRK2_S148F_ with GIRK1, GIRK1*, or the GIRK1_ΔdCT_ variants, with or without Gβγ, had small currents and were not blocked by ETX or by Ba^2+^ (Fig. S10B,F,H,I). Chen et al. (Chen et al., 2022) found that the homotetrameric GIRK2_S148F_ is a distorted channel, non-selective to K^+^ and devoid of Ba^2+^ block, due to the formation of ion conduction pathways besides the SF. Consequently, lack of ETX or Ba^2+^ block cannot be taken as an indication of loss of their binding to the channel.

When we analyzed the current-voltage relationships of GIRK1/2, we noticed that the apparent affinity for ETX was moderately voltage dependent (Fig. S11). K_d,app_ decreased at depolarized membrane potentials, with or without Gβγ. In contrast, K_d,app_ of ETX in GIRK1*/2, GIRK1*/2+Gβγ, as well as in homotetrameric GIRK1* and mGIRK2 remained constant at all potentials. The voltage-dependence of GIRK1/x channels depends of F137 in GIRK1 (Kofuji et al., 1996); our findings further indicate an allosteric coupling between voltage-dependence and ETX sensitivity, which involves F137.

## Discussion

We investigated the mechanism of inhibition (block) of GIRK channels by ETX. We report that ETX block is allosteric, subunit-specific, and is enhanced by Gβγ. Focusing on GIRK2, the best-studied and highly ETX-sensitive subunit, we uncover the molecular and structural basis of ETX action and its allosteric regulation by Gβγ.

GIRK blockers act by diverse mechanisms. Ba^2+^ binds within the SF, while the peptide tertiapin acts from the extracellular side (Alagem et al., 2001; Jiang & MacKinnon, 2000; Jin et al., 1999; Li et al., 2016; Patel et al., 2020). Block by the non-sedating antihistamine fexofenadine shows some Gβγ-dependency, but the mechanism is unclear (Chen et al., 2019). The anesthetic halothane either inhibits or enhances GIRKs in a subunit- and Gβγ-dependent manner, with a potential involvement of the HBC gate (Styer et al., 2010; Weigl & Schreibmayer, 2001); however, the binding site and mechanisms of action and Gβγ-linkage are unknown.

We postulate that ETX inhibits GIRKs through direct binding, because the block occurs in the absence of activators (Fig. 2) and is abolished by mutations of the putative ETX binding site in GIRK2 (Figs. 5, 8). Gβγ increases the apparent affinity of ETX, in parallel with channel activation; this suggests state-dependent blocking, as supported by kinetic modeling (Fig. 3). We rule out block by competition with Gβγ, because in this case adding Gβγ would decrease the apparent affinity of ETX. We also rule out steric block by ETX binding within the pore, because i) the block persists in a constitutively opened channel, GIRK2_N94H_, or upon activation with ivermectin (Fig. 3); ii) MD and mutagenesis locate the ETX binding site outside the ion conduction pathway. We conclude that ETX inhibition of GIRK channels, and its coupling to Gβγ-induced activation, are allosteric.

ETX inhibition varies greatly with channel’s subunit composition. In the absence of Gβγ, GIRK1/3 and GIRK1/4 are least and GIRK1/2 and GIRK2 are most sensitive (Fig. 2). Gβγ activation amplifies this divergence: GIRK1/3 and GIRK1/4 remain ETX-insensitive, whereas GIRK1/2’s apparent affinity rises mildly (twofold) and GIRK2’s dramatically (>30-fold). These findings suggest differential binding of ETX to GIRK subunits, and subunit specificity in coupling between Gβγ activation and ETX block. Clearly, GIRK2 and the homologous GIRK4 bind ETX (Fig. 3E). The interpretation of GIRK1 and GIRK3 data, based on heterotetrameric channels, is less straightforward. However, the choice of GIRK3 as the non-binder of ETX was subsequently confirmed in GIRK2-GIRK3 domain- and residue-swapping experiments (Figs. 5, 8). Of note, the significant difference in ETX block of GIRK1/2 and GIRK1/4 (with or without Gβγ) suggests that the presence of a Gβγ-ETX-sensitive subunit per se does not ensure ETX block and its Gβγ-induced enhancement. The conformation of a GIRK subunit probably varies within homo- vs. heterotetramers (Ponstingl et al., 2005), which may alter the affinity of ETX binding or the ETX-induced allosteric change.

GIRK1 initially also appeared as a non-ETX binder. However, the high ETX sensitivity of homotetrameric GIRK1* (Fig. 9), where F137 is exchanged to serine (conserved in other GIRK subunits), suggests otherwise. A straightforward interpretation of this result would be to assume that S148 of GIRK2 is an important part of the ETX-binding site; however, structural considerations (Fig. 10), as well as the presence of this serine in GIRK3, do not support this assumption. We conclude that F137 of GIRK1 and S148 of GIRK2 affect ETX sensitivity to ETX by an allosteric mechanism.

We located the ETX binding site using MD and mutagenesis. A TMD location was indicated by the domain swapping between GIRK2 and GIRK3 (Fig. 5). Unbiased flooding MD simulations on mGIRK2 suggested that ETX binds to a single subunit, close to the activator PIP_2_ (Fig. 6). We validated these predictions experimentally: swapping the putative ETX-binding region in GIRK2 (^90^KWRFN^94^ Fig. 8A) with GIRK3’s QWRLS fully or partially eliminated the ETX sensitivity in GIRK2 or GIRK1/2, and the inverse swap increased ETX block of GIRK1/3.

The MD analysis also offers critical insights into the mechanism of the allosteric block by ETX. Binding of ETX displaces PIP_2_ from its original binding site, and leads to a narrowing of the HBC gate (Fig. 6, Fig. S6). Shortest pathway analysis identifies important changes in the residue interaction network upon ETX inhibition. Highly conserved residues W91 and K194, as well as L89, K90 and F93 (non-conserved) were identified as critical (Fig. 7). Furthermore, a key role of N94 for information transfer in the allosteric gating pathway is uncovered by the network centrality analysis (Fig. 10). In support, mutations of N94 in GIRK2 to serine (as in GIRK3; Fig. 8) or histidine (N94H; Fig. 3) greatly weaken (though not fully abolish) the ETX block. Notably, the Gβγ-induced shift in ETX affinity in GIRK2_N94H_ and GIRK1/GIRK2_N94H/S_ channels is completely absent. These findings reveal a role for N94 as part of mechanisms underlying both the Gβγ-triggered allosteric change in GIRK2, and the coupling between Gβγ gating and ETX block. The unique role of N94 is supported by the fact that a lysine at the homologous position (K80) in ROMK1 is responsible for pH-dependent conformational changes that induce pore closure (Rapedius et al., 2007).

Interestingly, although both N94H and N94S mutants lose sensitivity to ETX, their gating properties are different. GIRK2_N94H_ has high basal activity and a reduced Gβγ activation (Yi et al., 2001) (Fig. 3), whereas GIRK2_N94S_ and GIRK1/2_N94S_ show basal and Gβγ-activated currents like the WT controls (Fig. 8, Fig. S7). Crystal structures (Whorton & MacKinnon, 2011, 2013; Yi et al., 2001) reveal that N94 is located in TM1, behind the F192 of the HBC gate located in TM2. The difference between N94H and N94S may arise from the larger volume of histidine that contains an imidazole ring, which might disrupt the HBC gate, stabilizing the open conformation. Importantly, for GIRK2_N94S_, the ETX block emerges as a sensitive indicator of conformational changes undetected by the standard parameters of channel gating (I_basal_ and Gβγ-induced activation).

S148 of GIRK2, strategically located behind the SF, emerges as another important component in the ETX-Gβγ-gating axis (Fig. 10). The corresponding F137 in GIRK1 is a well-known gating regulator. It controls a voltage-dependent aspect of GIRK1 gating (Kofuji et al., 1996), single channel conductance (Chan et al., 1996), and, as shown here, open probability and selectivity (Figs. S9, S11). Our finding of voltage sensitivity of ETX block in GIRK1/2, and its loss after the mutation of F137 to serine (Fig. S11) is in line with the allosteric linkage between ETX block and S148. Overall, the standard macroscopic parameters of GIRK1* and GIRK1*/2 (I_basal_ and Gβγ activation) are indistinguishable from other typical GIRK1/x channels (Fig. 8, Fig. S10); the dramatic change in ETX action again highlights its exquisite sensitivity to conformational changes in GIRKs. Taken together, our findings indicate that the inability of GIRK1’s F137 to propagate the allosteric change in the ETX-Gβγ-gating axis, and not lack of ETX binding, is the sole reason for the low ETX sensitivity of GIRK1/x channels and for the emergence of high ETX sensitivity in GIRK1*.

We further propose that, in GIRK1/2, the Gβγ-induced increase in ETX potency results from a Gβγ-induced allosteric change in GIRK2 subunits. The effect of Gβ_1_ mutants Gβ_(I80T)_ and Gβ_(I80N)_ reinforces this hypothesis. These Gβγ mutants are LoF for GIRK2, but activate GIRK1/2 normally, probably mainly via GIRK1 (Reddy et al., 2021). However, the shift in apparent affinity of ETX is lost (Fig. 4). Once again, ETX apparent affinity is sensitive to otherwise undetectable nuances in channel activation, in this case for Gβ_I80T/N_γ- vs. Gβγ_WT_- activated GIRK1/2.

From a physiological point of view, our finding of an increased ETX potency after Gβγ activation underscores the potential role of GIRKs as ETX targets in the treatment of epilepsy. After Gβγ activation, the sensitivity to ETX of the main neuronal GIRK, GIRK1/2 (K_d,app_ ∼1 mM), is significantly higher compared to VGCCs (K_d,app_ >10 mM) in our model system. Homotetrameric GIRK2 channels are abundant in the midbrain, but may be present in other parts of the brain along with GIRK1/2 (Lujan & Aguado, 2015; Luscher & Slesinger, 2010). The Gβγ-activated GIRK2 is exquisitely ETX-sensitive (K_d,app_ ∼0.1 mM). After Gβγ activation, the extent of inhibition of these neuronal GIRKs by therapeutic doses of ETX is 30-80%. It sounds counterintuitive to think of a K^+^ channel blocker as an antiepileptic; usually this is expected from K^+^ channel openers (Jeremic et al., 2021). However, as discussed previously (Colombo et al., 2023), inhibition can be beneficial if the epilepsy arises from K^+^ channel hyperactivity in inhibitory neurons, or when the depolarization caused by the inhibition of K^+^ channels inactivates T-type VGCCs that are often involved in the initiation and propagation of epileptic activity (Cheong & Shin, 2014; Rajakulendran & Hanna, 2016). For drug development purposes, allosteric modulators are considered superior to pore blockers, which often block a variety of similar channels. Allosteric modulators, which interact with channels in non-pore domains, are more specific (Zhang et al., 2020). Thus, ETX may become a prototype for future development of higher-affinity, specific allosteric blockers or even openers of GIRKs.

## Materials and methods

### Ethical approval of Xenopus laevis, oocyte preparation, and electrophysiology

Experiments were approved by Tel Aviv University Institutional Animal Care and Use Committee (permit #01-20-083). Maintenance and surgery of female frogs were as described previously (Rubinstein et al., 2009). In brief, adult female *Xenopus laevis* frogs were purchased from Xenopus 1 (Dexter, MI). For surgery, frogs were anesthetized in 0.2% tricaine methanesulfonate (MS-222). After collection of oocytes from the ovary, frogs were stitched and allowed recovery from anesthesia. Then, frogs were returned to a separate tank for post operational animals. Oocytes were defolliculated in Ca^2+^ free solution (in mM: 96 NaCl, 2 KCl, 1 MgCl_2_, 5 HEPES, pH 7.5) with collagenase. Two hours later, oocytes were washed with NDE solution (in mM: 96 NaCl, 2 KCl, 1 MgCl_2_, 1 CaCl_2_, 5 HEPES, 2.5 mM sodium pyruvate, 50 mg/ml gentamycin, pH 7.5 (Rubinstein et al., 2007)) and let to rest for at least 2 hours before injection. Then, oocytes were injected with RNA and incubated for 3 days at 20–22◦C in NDE solution.

### Electrophysiology

For electrophysiological experiments, standard two electrode voltage clamp (TEVC) setup was used. Oocytes were placed in ND96 solution (low K^+^) (in mM: 96 NaCl, 2 KCl, 1 MgCl_2_, 1 CaCl_2_, 5 HEPES, pH 7.5) or directly in high 24 mM K^+^ solution (HK24; in mM: 24 KCl, 74 NaCl, 1 MgCl_2_, 1 CaCl_2_, 5 HEPES, pH adjusted to 7.5 with KOH) at 20–22◦C. Whole-cell GIRK currents were measured using GeneClamp 500B amplifier, and Axon Digidata 1440a (Molecular Devices). Ethosuximide (Sigma-Aldrich, E7138) and the pore blocker Ba^2+^ (Merck, 101719) containing solutions were prepared freshly for each experiment. Ethosuximide was dissolved in HK24 solution to highest concentration. The next concentration was made by diluting the higher dose sequentially, until the smallest dose. Pressurized perfusion system (AutoMate Scientific) was used for solution exchange. 2 µM Ivermectin was used to activate GIRK2 channels (Alomone labs, I-110). Experiments were performed as follows: after current was stabilized in ND96 or HK24 solutions, 2 seconds voltage ramp from -120 mV to +50 mV was applied to obtain I-V curves. Then the solution was exchanged to the HK24 with the lowest ETX concentration, after current stabilization another voltage ramp was applied. All ETX concentrations were applied sequentially, the recording was finished with application of HK24 with 2.5 mM Ba^2+^, Ba^2+^ was used to subtract endogenous non GIRK currents.

Recording of VGCCs was performed in high-Ba^2+^ solution, in mM: 40 Ba(OH)_2_, 50 mM NaOH, 2 mM KOH, and 5 mM HEPES, titrated to pH 7.5 with methanesulfonic acid (Oz et al., 2011). Oocytes were held at - 60 mV. Peak currents were measured by voltage steps from -80 mV to -10 or +20 mV for Ca_V_3.2 and Ca_V_1.2, respectively. Leak subtraction was done off-line; leak conductance was calculated from a voltage step from - 80 to -100 mV.

Patch clamp experiments were done using Axopatch 200B as described (Yakubovich et al., 2015). Currents were recorded in cell-attached mode, at -80 mV, filtered at 2 kHz and sampled at 20 kHz. Patch pipettes had resistances of 1.4–3.5 MΩ. Pipette solution contained, in mM: 146 KCl, 2 NaCl, 1 MgCl_2_, 1 CaCl_2_, 1 GdCl_3_, 10 HEPES/KOH (pH 7.5), with or without 2 µM acetylcholine (ACh). GdCl_3_ completely inhibited the stretch-activated channels. The bath solution contained, in mM: 140 KCl, 6 NaCl, 2 MgCl_2_, 1 EGTA, 10 HEPES/KOH (pH 7.5).

### DNA constructs and RNA

cDNA constructs of rat GIRK1, rat GIRK1_ΔdCT_, human GIRK1*, rat GIRK1*_ΔdCT_, mouse GIRK2A, human GIRK2, rat GIRK3, and rat GIRK4, bovine Gβ_1_ (WT, K78R, I80N, I80T), and bovine Gγ_2_ were all inserted into the pGEM-HJ vector containing 5’ and 3’ untranslated regions from *Xenopus* β-globin (Kahanovitch et al., 2014; Rubinstein et al., 2009; Shalomov et al., 2022; Tabak et al., 2019). Human GIRK2, with modified sequence for better expression in Chinese Hamster Cells (CHO), in pcDNA3.1 was obtained from Blavtnik center for drug discovery in Tel Aviv University. Human GIRK2 was subcloned into pGEM-HJ (Table 3) using GibON kit (ERX-E1050, Eurx). DNAs of VGCCs: rabbit α1C, rabbit α2δ, rabbit β2b, human α1H, and rat Cachd1 (Table S5). Chimeras of GIRK2 TMD and GIRK3 CSD, and GIRK3 TMD and GIRK2 CSD were also prepared using GibON kit (ERX-E1050, Eurx). Triple or point mutations were done in mGIRK2 and rGIRK3: GIRK2_QLS_, GIRK2_K90Q_, GIRK2_F93L_, GIRK2_N94H_, GIRK2_N94S_, GIRK3_KFN_, GIRK3_Y350A_. All primers for DNA manipulation are presented in Table S4. mRNAs were prepared as described previously (Dascal & Lotan, 1992). The amounts of mRNA injected per oocyte were varied according to the experimental design and presented in the text or figures. For maximal channel activation by Gβγ, we injected 5 ng Gβ and 1 ng Gγ mRNA. These weight ratios were chosen to keep approximately equal molar amounts of Gβ and Gγ RNA.

### Modeling of Gβγ and ETX interaction with GIRK2

To simulate GIRK2 activation by Gβγ, we utilized cooperative sequential model proposed by Wang et al. (Wang et al., 2016), that assumes that the channel binds 1 to 4 Gβγ molecules, and only 4-Gβγ occupied channel is available for opening (Fig. 3D). Na^+^ was kept constant during the whole time of experiment and the level of PIP_2_ was assumed to remain constant; thus we utilized single constant cooperativity factor b=0.3 (Wang et al., 2016). Channel activity was calculated according to Klotz-like cooperativity scheme (Klotz, 2004). To model ETX interaction with GIRK2, we tested several inhibition schemes (see sup. Fig.1 and sup. Table 1). For simulation, we constructed systems of differential equations based on these schemes and solved them numerically in Berkley Madonna (Berkeley Madonna, Inc., Albany, CA) (Marcoline et al., 2022) utilizing 4th order Runge-Kutta integration method. The simulations were run till apparent steady-state (see Supplementary Material for description). To estimate the opening from C_4_ (α) and closing (β) rate constants of GIRK2 channel, we performed a limited single channel analysis of GIRK2 activity using Clampfit 10 (Molecular Devices, San Jose, USA), as detailed in Supplementary Methods.

### Molecular dynamics simulations

An open, conductive conformation of the Kir3.2 channel (PDB code: 3SYA) (Whorton & MacKinnon, 2011) in a Berger lipid POPC membrane (Berger et al., 1997) bound to the activator PIP_2_ was extracted from a previous MD simulation (Bernsteiner et al., 2019). Ten copies of each R-ETX and S-ETX were placed randomly in the simulation box, resulting in a concentration of 18.2 mM. ETX was parametrized based on GAFF2 (Wang et al., 2004). The simulations were carried out as described previously (Friesacher et al., 2022) using GROMAX5.1.2 (Abraham et al., 2015) in combination with amber99SB (Hornak et al., 2006). Five replicas of the system were simulated for a total of 1.5 μs. Analysis of distances, densities, hydrogen bonds and RMSDs were carried out using the Gromacs analysis tool box. Computation of shortest paths, betweenness centrality and closeness centrality were based on the protein energy network analysis performed with the “get Residue Interaction Energies and Networks” (gRINN) tool (Sercinoglu & Ozbek, 2018). PyMol 2.7. (The PyMOL Molecular Graphics System, Version 2.3.0 Schrödinger, LLC.) was used for the calculation of the solvent accessibility and visualization of the system.

### Data analysis and statistics

Analysis of electrophysiological recordings was performed with pCLAMP (Molecular Devices) and custom software written for this work using Python 3.10 and the pyABF package (https://pypi.org/project/pyabf). Single channel current (i*_single_*) was calculated from all-point histograms of the original records, and open probability (P_o_) was obtained from event lists generated using idealization procedure based on 50% crossing criterion, as described (Yakubovich et al., 2015). Number of channels was estimated from overlaps of openings during the whole time of recording (at least 4 min). Both mean open time (t_o,mean_) and open probability (P_o_) was calculated only from records that contained 1 or 2 channels. Additional details are described in Supplementary Methods.

In ETX dose-response analysis (the two-electrode voltage clamp experiments), we excluded from analysis records with clear mechanical disturbances caused by perfusion. Current amplitudes at -80 mV were extracted from the I-V relationships recorded in all HK24 solutions, with all ETX concentrations and with Ba^2+^. Net GIRK currents were obtained by subtraction of current recorded in HK24 with Ba^2+^. Percent of inhibition was calculated as follows:

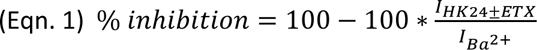

For ETX dose-response curves, GraphPad Prism for Windows (GraphPad Software, Boston, Massachusetts USA) was used to fit the data to Hill equation (Eqn. 2) or two component equation (Eqn. 3) for dose response with IVM activated GIRK2:

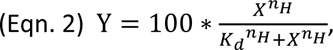

where Y is % of inhibition by ETX. K_d,app_ and Hill coefficient (n_H_) were extracted from the fit and used for further analysis.

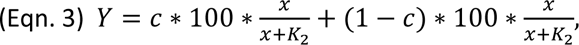

where *K_1_* and *K_2_* are the ETX dissociation constants from the high- and low-affinity sites, respectively, and c is the fraction of the higher affinity population corresponding to K_1_.

Statistical analysis was done using GraphPad Prism. Outlier values of n_H_ and K_d_ were identified by ROUT method and excluded. Data in the figures are presented as mean ± SEM for normally distributed data, and otherwise as box with median and min-max whiskers. Statistical tests were: unpaired t-test or One-way ANOVA with Tukey post hoc test for distributions that passed Shapiro-Wilk normality test, otherwise Mann-Whitney or Kruskal-Wallis with Dunn’s post hoc test were used.

## Author contributions

Conceptualization: ND, ASW, BS, TF

Experiments design: ND and BS

Experiments performance: BS, HPR, JCC, SD

Analysis of experiments: B, ND, HPR, JCC

Kinetic modeling: DY

Computational design: ASW, TF, EMZP

Molecular modeling and analysis: TF, HB, ASW, TF, EMZP

Writing – original draft: BS, TF, DY

Writing – review and editing: ND, ASW, EMZP, BS, TF, DY

All authors read and approved the paper

## Data and materials availability

Data and materials (mutant construct of GIRK subunits) will be made available upon reasonable request for the corresponding authors. Results of electrophysiology experiments were analyzed using the Clampfit 10.7 software (Molecular Devices). The custom software written for this work using Python 3.10 and the pyABF package (https://pypi.org/project/pyabf) for the analysis of GIRK I-V relationships is available upon request to Boris Shalomov. Visualization and analysis were performed using VMD, Pymol, Python, Java and the FMA tool. All of these software packages are publicly available.

## Supplementary Materials

### Supplementary Text

#### Kinetic modeling of Gβγ and ETX interaction with GIRK2

##### Single channel properties of GIRK2

We conducted limited kinetic analysis of mGIRK2 activation by saturating concentration of Gβγ. We analyzed cell-attached patches in oocytes injected with 5 ng RNA of Gβγ. Experiments in which one or two channels were observed (according to number of opening overlaps) were included in the analysis. The recordings were idealized utilizing 50% crossing criterion (Sakmann & Neher, 1995) and histograms of open times were constructed. Subsequently, form six attached patch recordings with one of two channels (exemplified in Supplementary Fig. 3A), we estimated the open probability (P_o_) to be 0.096±0.025 (mean±SEM, n=6) and the mean open time, t_o,mean_ = 1±0.34 ms (n=6), by fitting the open times distribution with a single exponent (Supplementary Fig. 3B). Since 5 ng Gβγ RNA activated the channel almost fully, we assumed that the observed P_o_ corresponds to the maximal P_o_, P_o,max_. For calculations, we used P_o,max_=0.1. Based on these data, for the transition C_4_⟺O, the opening (α) and closing (β) rate constants of GIRK2 channel could be calculated based on:

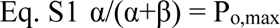

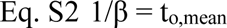

##### Calculation of kinetic constants for the schemes shown in Supplementary Fig. 3

From here, the forward and backward rates for the C_4_⟺O transition were: α=111.1 s^-1^; β=1000 s^-1^. We next simulated the block of GIRK2 by ETX, in the presence of increasing Gβγ concentrations, for schemes 2-5 of the Supplementary Fig. 3C. The simulations of the ETX block of GIRK2 for the allosteric block scheme (Fig. 3D, blue rectangle; Supplementary Fig. 3C, scheme #3) were done as follows. Based on principle of detailed balance (Jackson, 2006), the following relationships between the dissociation coefficients (Supplementary Fig. 3C, scheme #3) hold true:

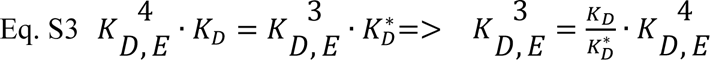

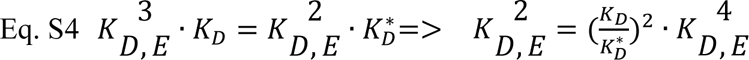

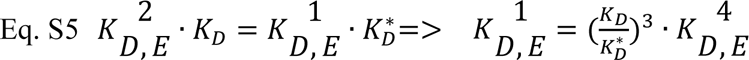

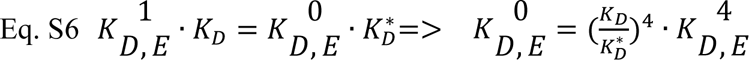

Here, K_D,E_ is the dissociation constant of ETX, K_D_ is the dissociation constant of Gβγ, * denotes Gβγ-bound channel. The superscripts 1 through 4 relate to the number of bound Gβγ molecules. For simplicity we assumed that 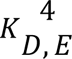 (the affinity of ETX to channel occupied by four Gβγ) was equal to IC_50_ of ETX obtained from the fit of dose-response to ETX measured in cell injected with a high concentration of Gβγ (5 ng/oocyte RNA of Gβ). Combining Eq. S6, 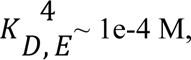 we tested a range of 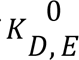 in order to obtain the ETX IC_50_ of ∼4 mM which was measured in oocytes injected with 0.2 ng/oocyte RNA of Gβγ (Fig. 3B, C), a very low concentration of Gβγ. We assumed that the high Gβγ concentration is of ∼ 100 molecules/µm^2^ while the low Gβγ concentration is of ∼ 2 molecules/µm^2^ (estimated from published data (Yakubovich et al., 2015)). We utilized the above mentioned dissociation coefficients in order to simulate the system of reactions of the schemes shown in Supplementary Fig. 3C. We implied Berkley Madonna in order to solve system of differential equations generated based on this scheme. The system was allowed to run till no visually observable change in reactant concentrations was seen, corresponding approximately to steady state values. Rate constants used for the simulations according to the allosteric block model are summarized in Supplementary Table 1C. All simulations were done for channel density of 10 channels/µm^2^. Microscopic reversibility rule was strictly implemented in all adjustments of the rate constants.

A similar approach was utilized for the case of open blocker channel scheme (Fig. 3D, dark red rectangle; Supplementary Fig. 3C, scheme #2). In particular, we tested a range of K_D,E_ values (10^-6^ to 10^-4^ M) and selected the K_D,E_ = 1e-5 M as rendering the closest to experimental IC50 values for low and high densities of Gβγ. Rate constants used for simulation of open channel block scheme are summarized in Supplementary Table 1B.

Another tested scheme was a closed channel blocker scheme (Supplementary Fig. 3C, scheme #4), where we assumed that ETX binds specifically to the C_4_ state and prevents the channel from entering the open state. Rate constants utilized for simulation are summarized in Supplementary Table 1D.

The last tested scheme was a closed block scheme with equal affinity i.e. it is assumed that ETC binds to all closed states with same affinity and prevents the transition from C_4_ to open state. This scheme is shown in Supplementary Fig. 3C (scheme 5). Rate constants and equilibrium constants utilized for simulation are summarized in Supplementary Table 1E. It can be seen from Supplementary Fig. 3E that, according to this scheme, no leftward shift of dose-response to ETX upon increase in Gβγ expression is expected.

## Supplementary Figures

**Fig. S1.**
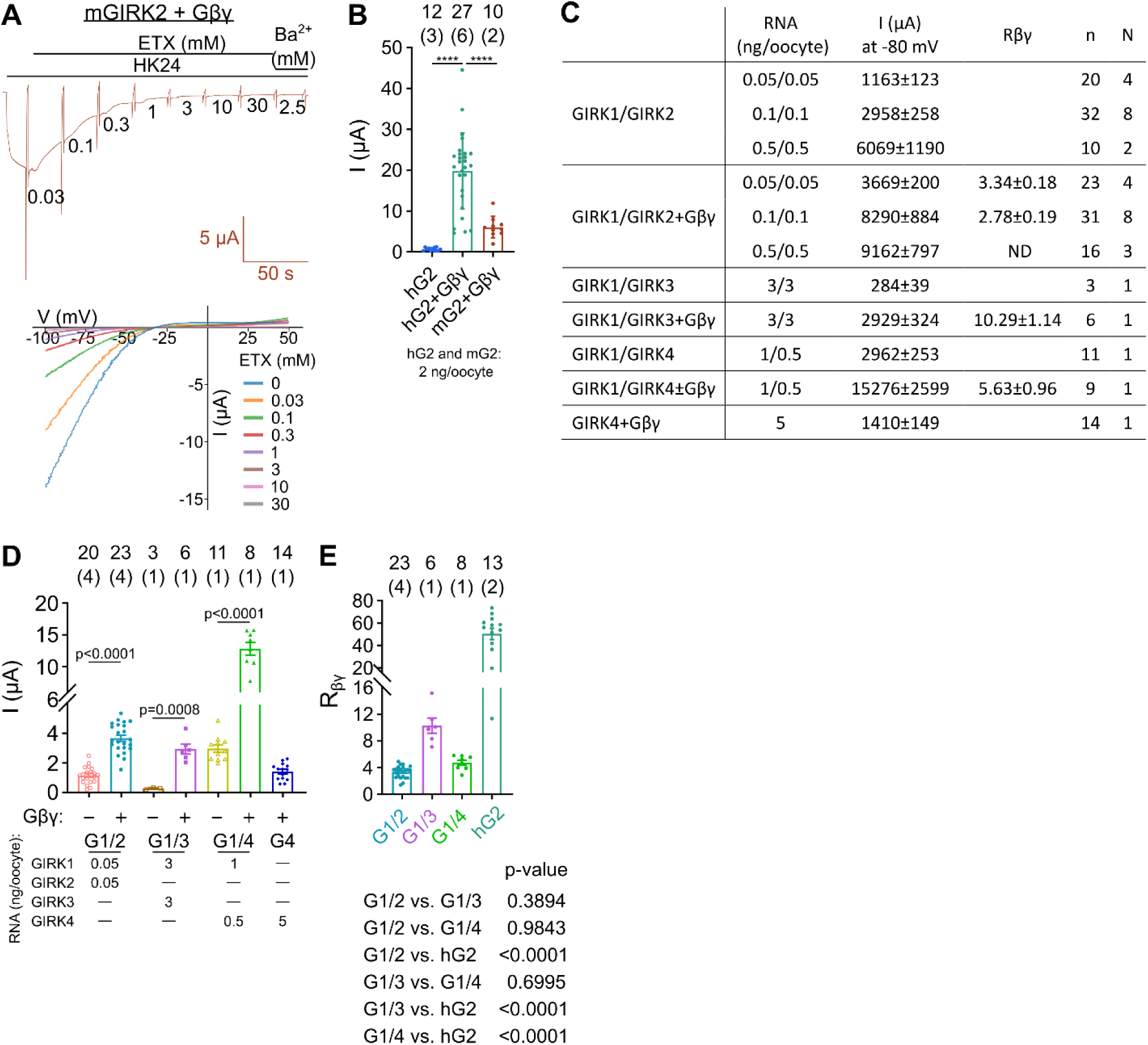
Current amplitudes of different subunit combinations of GIRK channels. In all cases where Gβγ was present, 5 ng Gβ and 1 ng Gγ RNAs were injected per oocyte. G1 through G4 stands for GIRK1 through GIRK4. **A**, an exemplary recording of the dose-dependent effect of ETX on mGIRK2 coexpressed with Gβγ (mGIRK2+Gβγ). **B**, current amplitudes of hGIRK2, hGIRK2+Gβγ, and mGIRK2+Gβγ. Numbers above bars denote number of cells (n), with number of experiments (N) in brackets. **** p<0.0001, one-way ANOVA followed by Tukey’s test. **C**, current amplitudes and fold activation by Gβγ (Rβγ; mean±SEM) of GIRK1/2, GIRK1/3, GIRK1/4 with and without Gβγ, and GIRK4+Gβγ. This is a summary of all experiments shown in Fig. 2. n, number of oocytes; N, number of experiments. **D**, comparison of basal and Gβγ-activated currents in a subset of experiments with the indicated amounts of channels’ RNA (bottom). Numbers above bars show number of cells with number of experiments (in brackets). RNA doses of GIRK subunits varied depending on their combination, thus only unpaired t-test was performed, to compare currents without and with Gβγ. **E**, Rβγ of GIRK1/2, GIRK1/3, GIRK1/4 and hGIRK2 from the experiments shown in **A** and **D**. Statistical comparison (bottom) was done by one-way ANOVA followed by Tukey’s test.

**Fig. S2.**
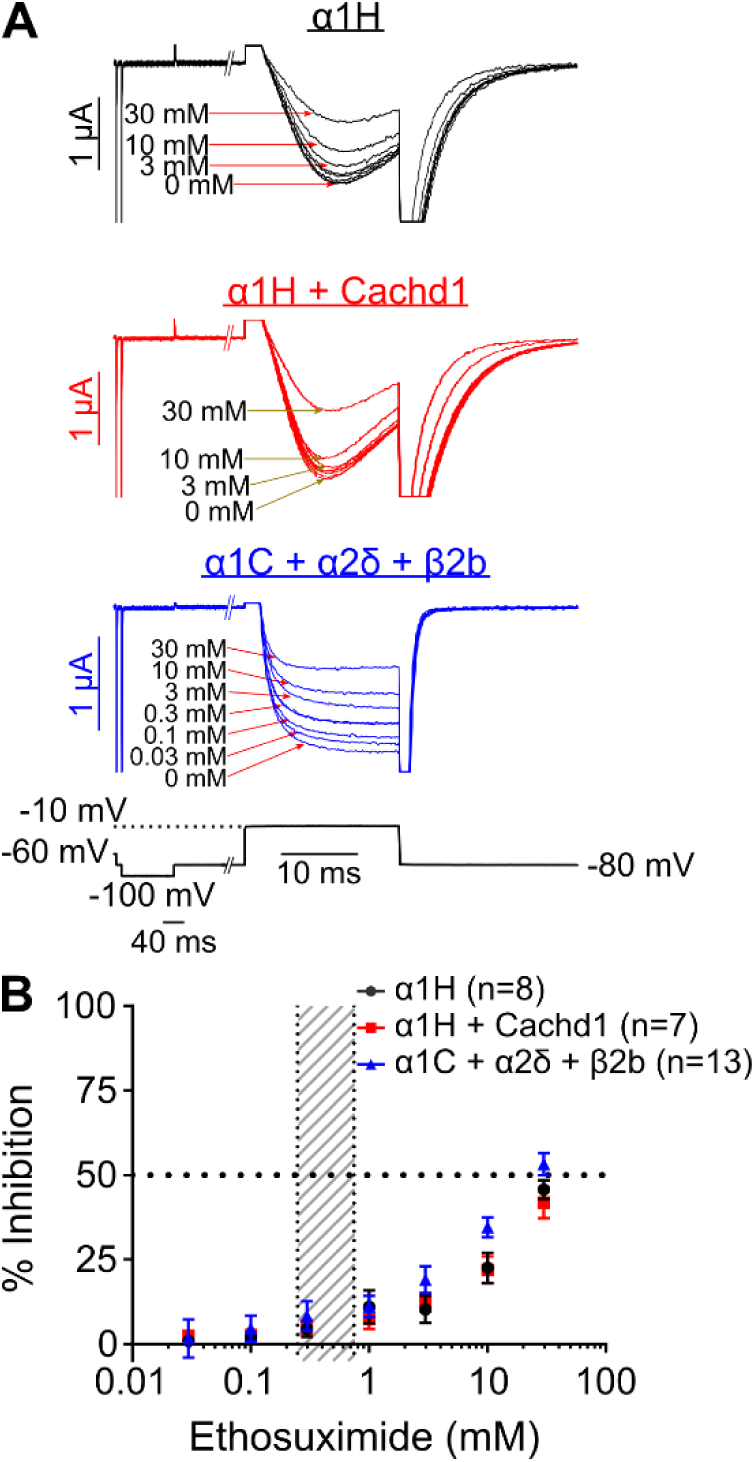
T-type and L-type Ca^2+^ channels are blocked by ETX with low affinity. **A**, examples of current records of α1H (Ca_V_3.2), α1H + a2δ like subunit, Cachd1, and α1C + a2δ + β2b (Ca_V_1.2), respectively, with increasing doses of ETX. Currents were measured by voltage steps from -80 mV to -10 mV for Ca_V_3.2, and from -80 to +20 mV for Ca_V_1.2, in a solution containing 40 mM Ba^2+^. **B**, dose dependence of ETX inhibition shows that the apparent affinity of ETX to T-type and L-type VGCCs is low.

**Fig. S3.**
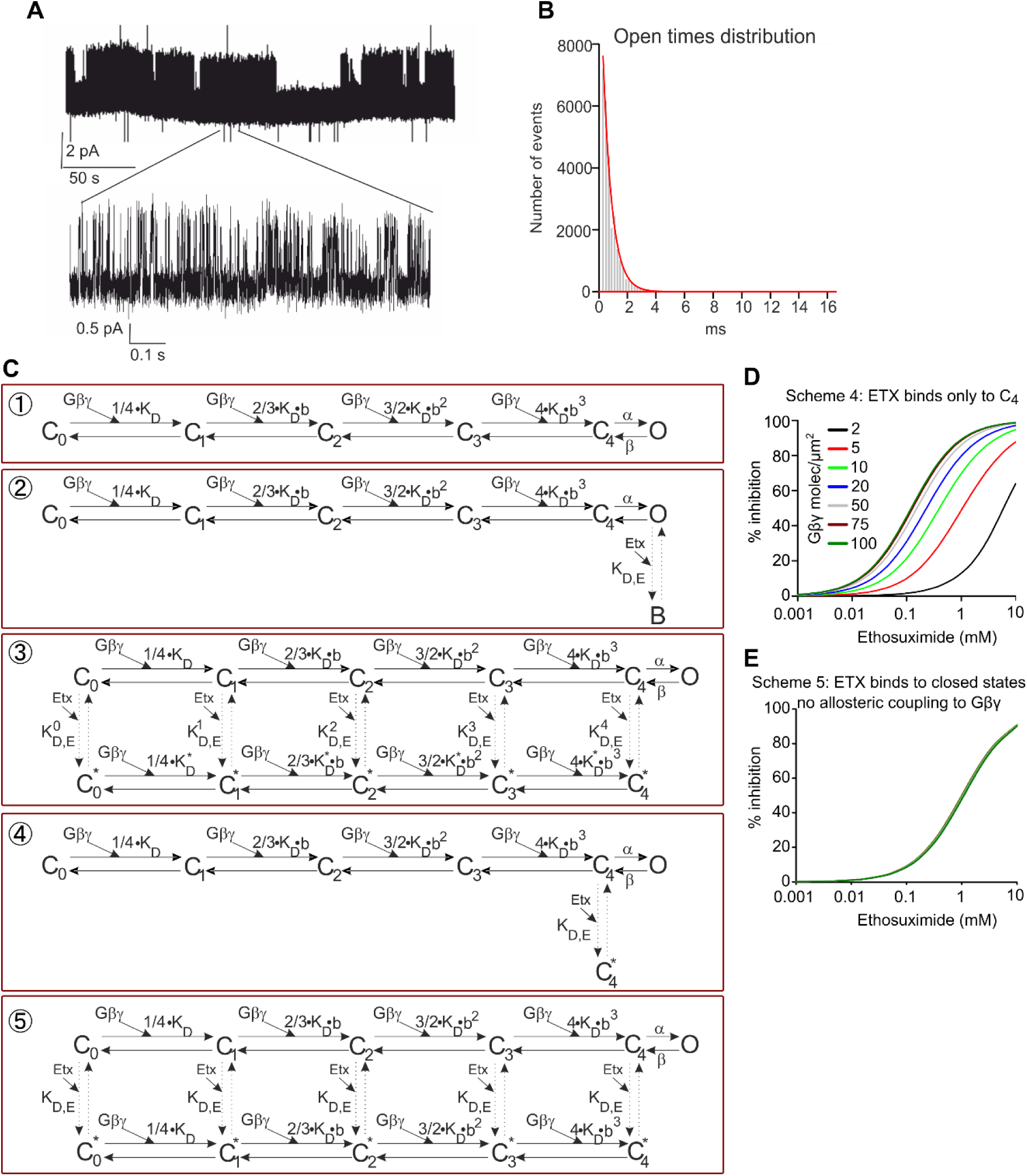
Kinetic modeling of ETX block of GIRK2. **A**, GIRK2 channel activity was recorded in oocytes injected with 5 ng/oocyte RNA of Gβγ. Openings are shown as upward deflections. **B**, open times histogram with a one-exponential fit (τ = 0.62 ms). **C**, the tested schemes of GIRK2-Gβγ-ETX interaction, explained in Supplementary Table 1. 1 – GIRK2 channel activation by Gβγ, 2 – open channel blocker scheme, 3 – allosteric closed channel blocker scheme, 4 – state-dependent closed channel blocker scheme (ETX interacts only with C_4_), 5 – state-independent closed channel blocker scheme (ETX binds to all closed sates with the same affinity and no assumption of change in ETX-occupied channel affinity to Gβγ is made). **D**, simulation of the dose-response to ETX with different level of expression of Gβγ utilizing scheme #4. **E**, simulation of the dose-response to ETX with different level of expression of Gβγ utilizing scheme #5.

**Fig. S4.**
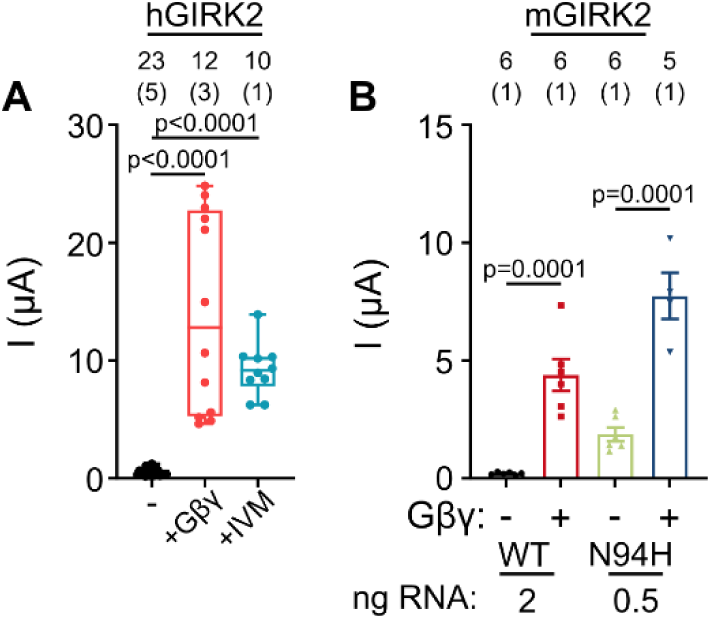
Effects of IVM and N94H mutation on GIRK2 current amplitudes. **A**, 2 µM IVM activates hGIRK2 (2 ng RNA) similarly to Gβγ (5 ng Gβ RNA, 1 ng Gγ RNA) (Kruskal-Wallis test followed by Dunn’s test). The difference between IVM- or Gβγ-activated current was not significant (p>0.999). **B**, mGIRK2_N94H_ has a high basal current and is additionally activated by Gβγ, while mGIRK2_WT_ has a small basal current and a much greater extent of activation by Gβγ than mGIRK2_N94H_ (unpaired t-test for all comparisons). Note that the high basal current was observed with only 0.5 ng RNA/oocyte of mGIRK2_N94H_, which is 4 fold less than for mGIRK2_WT_.

**Fig. S5.**
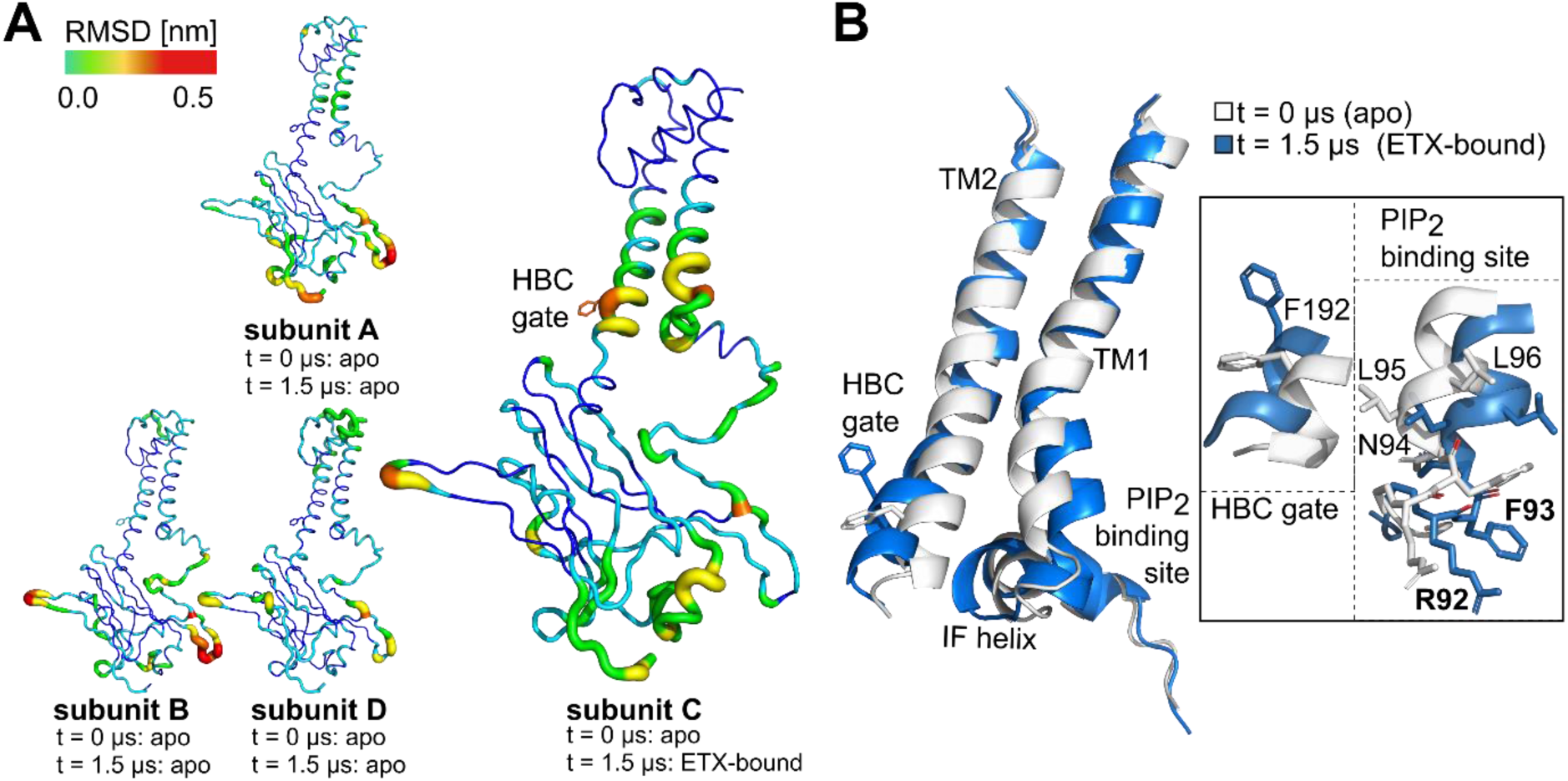
ETX binding-induced conformational changes of the GIRK channel observed in run1. **A**, GIRK2 subunits are colored based on an alignment between the subunit conformations at the beginning of the simulations (t=0 µs) and the end of the simulation (t=1.5 µs). The HBC gate is shown in stick representation. **B,** Alignment of the starting conformation (t= 0 µs, not bound to ETX; white) and the end conformation (t=1.5 µs, ETX-bound; blue) of subunit C. The HBC gate, the ETX-binding residues R92 and F93, and the residues in proximity to the ETX binding site are shown in stick representation. On the right, a close up of the alignment is shown, focusing on the ETX binding region at the TM1 and the HBC gating region.

**Fig. S6.**
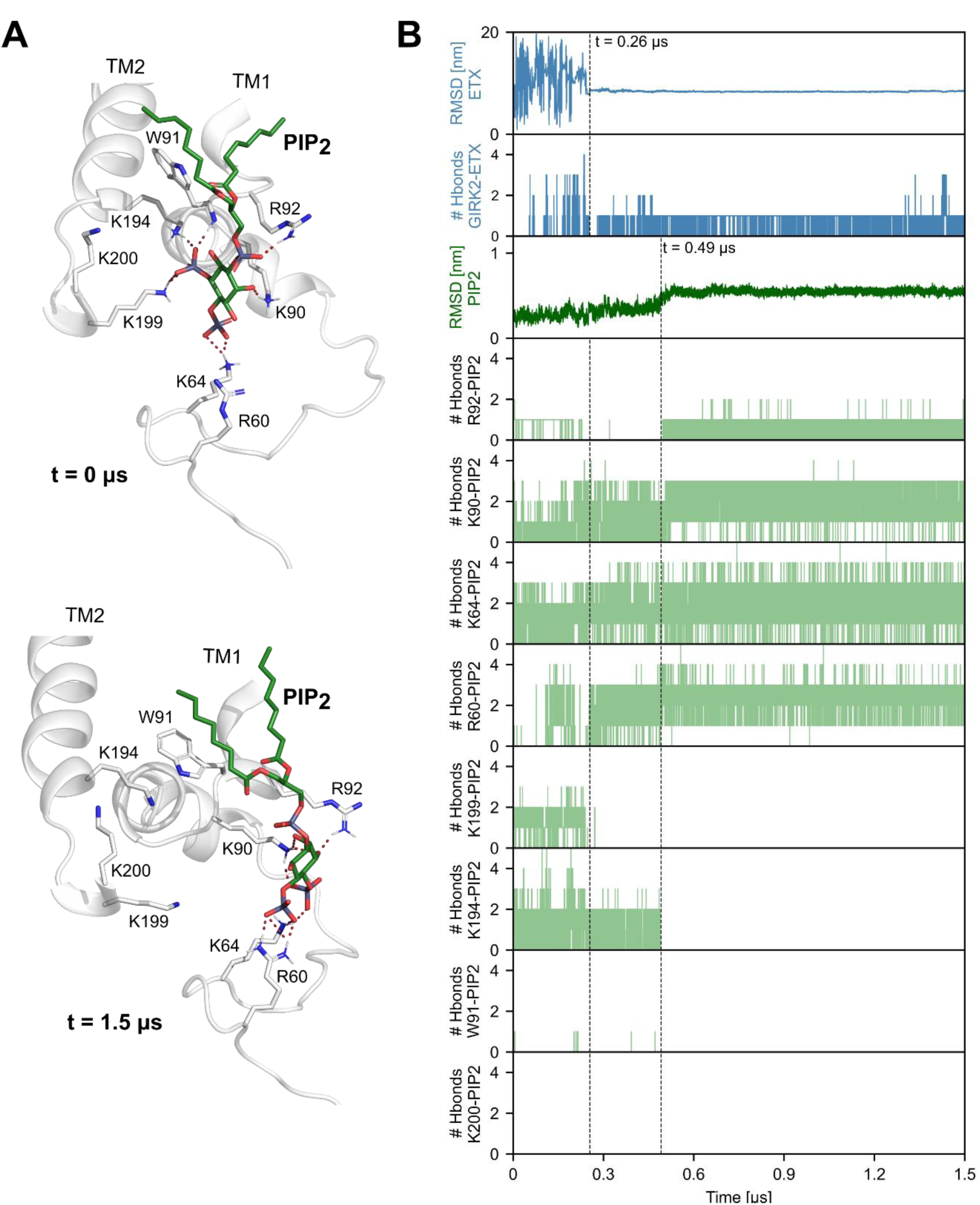
The effect of ETX binding on the PIP_2_ conformation in run1. **A,** the structures show the PIP_2_ conformation at the beginning (t= 0 μs) and at 1.5 μs of the MD simulation. PIP_2_ is shown in green and GIRK2 in grey. PIP_2_ and PIP_2_ binding residues of GIRK2 are shown as sticks. Red dashed lines represent hydrogen bonds. **B,** RMSDs and hydrogen bonds formation of ETX (blue) and PIP_2_ (green) over time. The dashed lines indicate the time ETX binds to GIRK2 (t= 0.26 µs), and where PIP_2_ is stabilized in a displaced conformation, 5.5 Å from its original position (t= 0.49 µs).

**Fig. S7.**
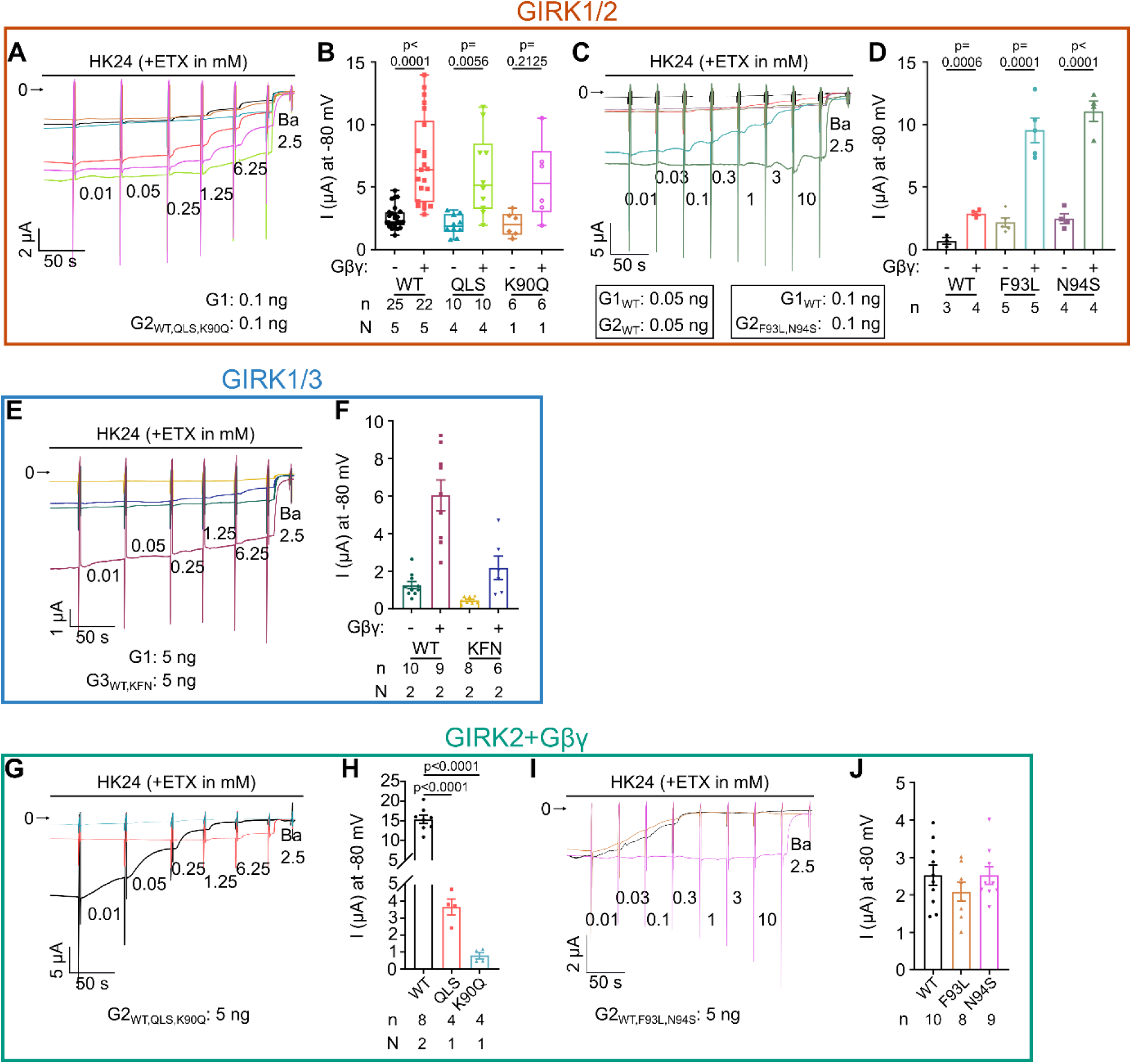
The effect of mutations in the PIP_2_ binding region. In all experiments the amount of injected RNA of Gβγ was the same (5 ng Gβ and 1 ng Gγ). The amount of RNA for channel in each experiment is noted in panels. **A**, representative original recordings (not normalized) of ETX effect on GIRK1/2_WT_, GIRK1/2_QLS_, and GIRK1/2_K90Q_. Color coding as in B. **B**, current amplitudes of GIRK1/2_QLS_ and GIRK1/2_K90Q_ with and without Gβγ were similar to GIRK1/2_WT_ (P>0.05; Kruskal-Wallis followed by Dunn’s multiple comparison test). **C**, current recordings of GIRK1/2_WT_, GIRK1/2_F93L_, and GIRK1/2_N94S_. Color coding as in D. **D**, current amplitudes of GIRK1/2_WT_, GIRK1/2_F93L_, and GIRK1/2_N94S_ (unpaired t-test for all comparisons). **E**, representative recordings of GIRK1/3_WT_, and GIRK1/3_KFN_. Color coding as in F. **F**, current amplitudes of GIRK1/3_WT_ and GIRK1/3_KFN_ (one-way ANOVA followed by Tukey’s multiple comparisons test). **G**, representative recordings of GIRK2_WT_, GIRK2_QLS_, and GIRK2_K90Q_. Color coding as in H. **H**, current amplitudes of GIRK1/2_WT_, GIRK2_QLS_, and GIRK2_K90Q_ (one-way ANOVA followed by Tukey’s multiple comparisons test). **I**, representative recordings of GIRK2_WT_, GIRK2_F93L_, and GIRK2_N94S_. Color coding as in J. **J**, current amplitudes of GIRK1/2_F93L_ and GIRK1/2_N94S_ are similar to GIRK1/2_WT_ (one-way ANOVA followed by Tukey’s multiple comparisons test).

**Fig. S8.**
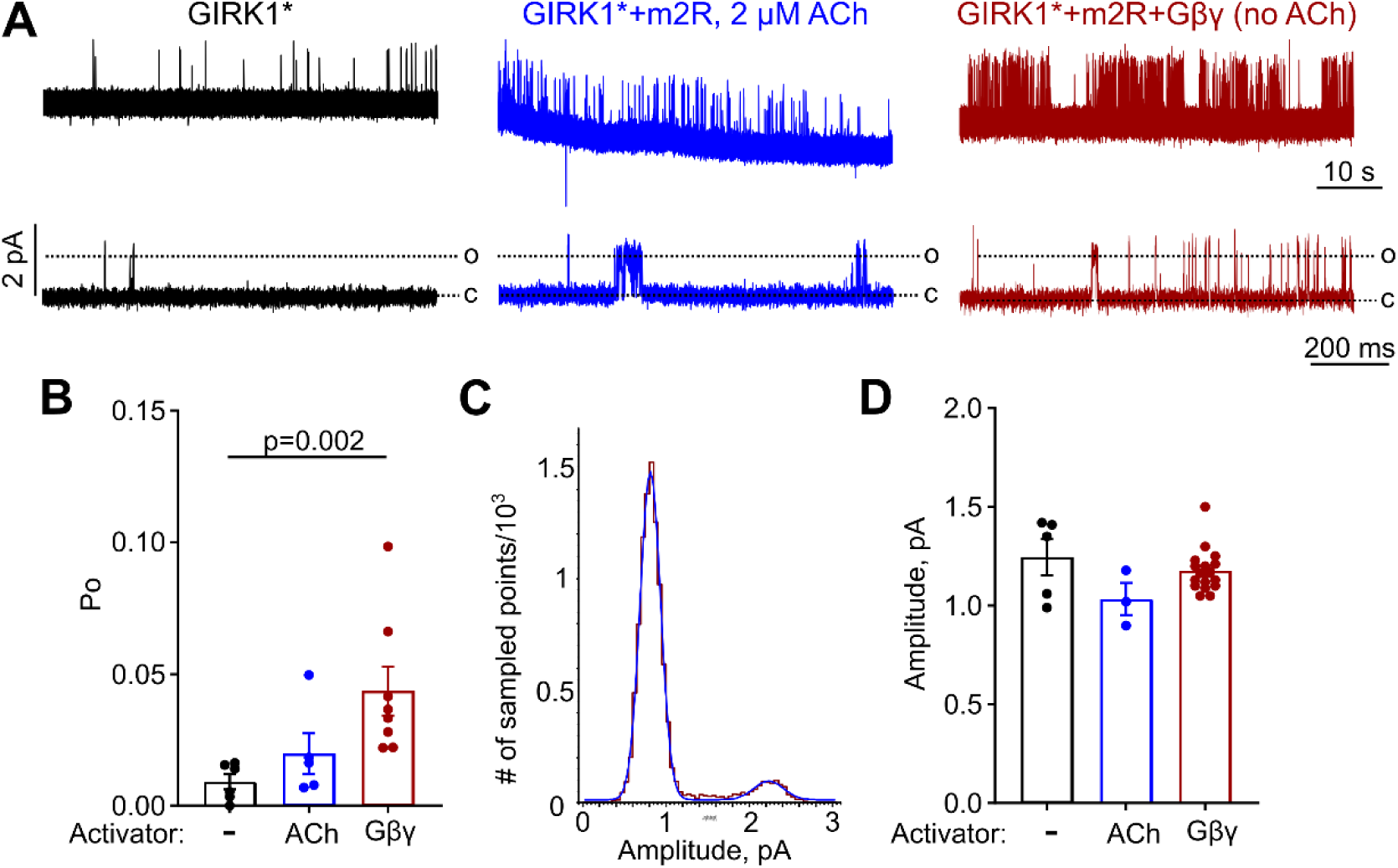
Single channel properties of GIRK1*. **A**, representative cell-attached, single channel recordings of GIRK1* expressed alone (left, in black); GIRK1* coexpressed with m2 receptor activated by acetylcholine (ACh) (middle, in blue; 1-5 ng m2 receptor RNA); and GIRK1* coexpressed with m2 receptor and Gβγ (right, in red; 5 ng Gβ and 1 or 2 ng Gγ RNA). c denotes the closed state and o denotes the open state of the channel. **B**, P_o_ of GIRK1*, GIRK1* activated by ACh, and GIRK1* activated by Gβγ. Gβγ had significantly increased P_o_ of GIRK1*. The average P_o_ of GIRK1* in the presence of Gβγ was 0.043±0.01 (n=8). **C**, a representative all-points histogram from a single channel recording in an oocyte expressing GIRK1* with Gβγ, used to assess the single channel amplitude, i_single_. Histogram (red line) was fitted to a two-component Gaussian (blue line). **D**, the single channel current amplitude of GIRK1* at -80 mV was similar in all conditions (p=0.21, Kruskal-Wallis test). The average (±SEM) i_single_ for GIRK1* in the presence of Gβγ, at -80 mV, was 1.18±0.03 pA (n=17). For comparison, for Gβγ-activated GIRK1/2 recorded under identical conditions, P_o_ is ∼0.105 and i_single_ is ∼2.8 pA (Yakubovich et al., 2015).

**Fig. S9.**
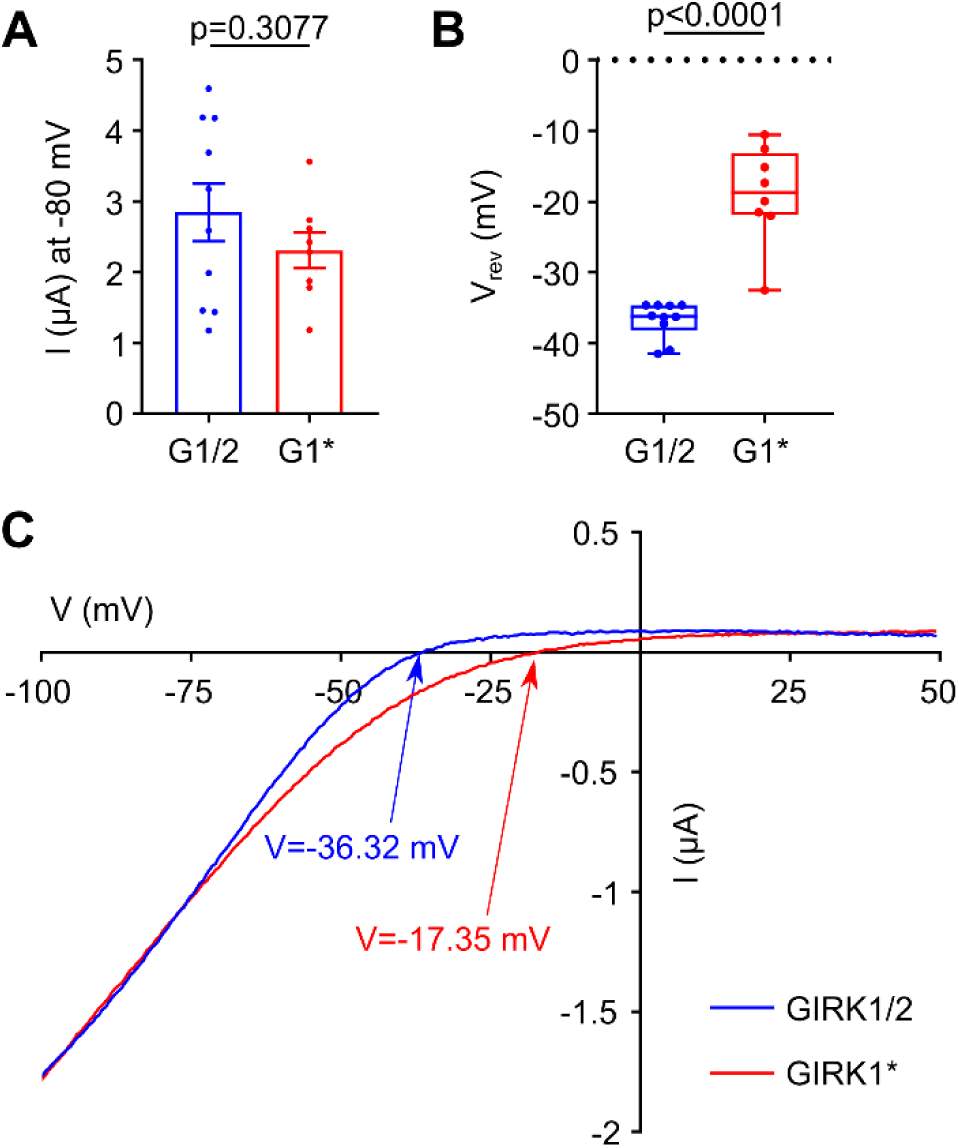
The reversal potential (V_rev_) of GIRK1* is depolarized compared to GIRK1/2. **A**, current amplitudes at -80 mV of GIRK1/2 and GIRK1* (n=10 and n=8, respectively). In this experiment, the channels were expressed with Gβγ. The RNA doses were (in ng/oocyte): GIRK1* 5, GIRK1 0.05, GIRK2-YFP 0.05, Gβ 5, Gγ 1. **B**, the reversal potential (V_rev_) in cells shown in A. V_rev_ of GIRK1* is more positive than of GIRK1/2_WT_ in HK24 solution. In HK24 solution, according to Nernst equation the expected E_K_ = -36 mV, assuming K_in_ = 100 mM (Dascal, 1987). **C**, I-V curves in representative oocytes.

**Fig. S10.**
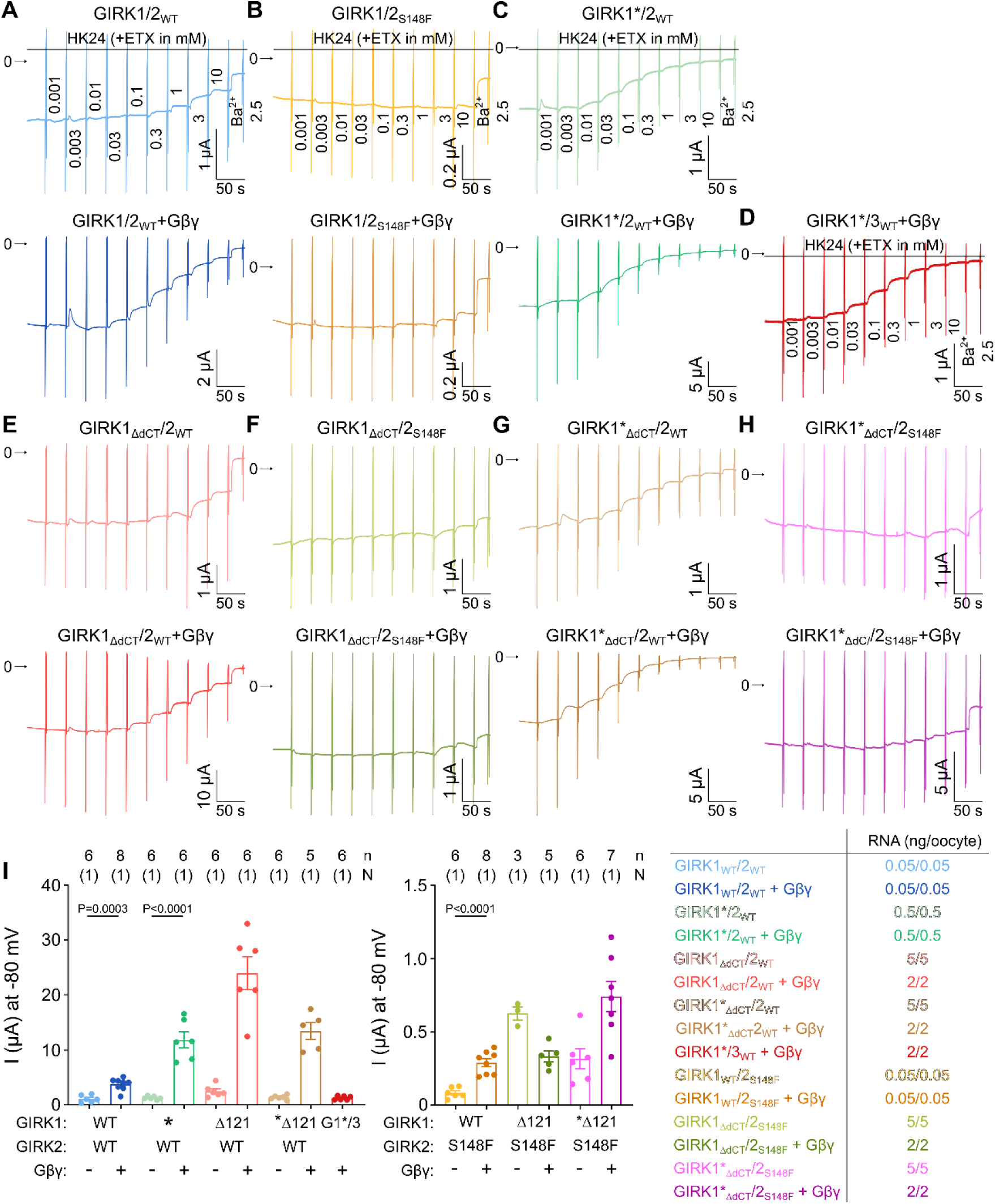
Representative current recordings and mean current amplitudes of GIRK1*- and GIRK2_S148F_-containing channels. Representative recordings without (top) and with Gβγ (bottom): **A**, GIRK1/2_WT_. **B**, GIRK1/2_S148F_. **C**, GIRK1*/2_WT_. **D**, GIRK1*/3_WT_+Gβγ. **E**, GIRK1_Δ121_/2_WT_. **F**, GIRK1_Δ121_/2_s148f_. **G**, GIRK1*_Δ121_/2_WT_. **H**, GIRK1*_Δ121_/2_s148f_. **I**, current amplitudes at -80 mV of the channels presented in **A-H** (left), and the amounts of injected RNA (right). Oocytes were injected with 5 ng of Gβ and 1 ng of Gγ in all cases. Note that the amounts of channel RNA without and with Gβγ were often different, therefore a direct comparison of amplitudes with/without Gβγ and an estimation of extent of Gβγ activation was not possible.

**Fig. S11.**
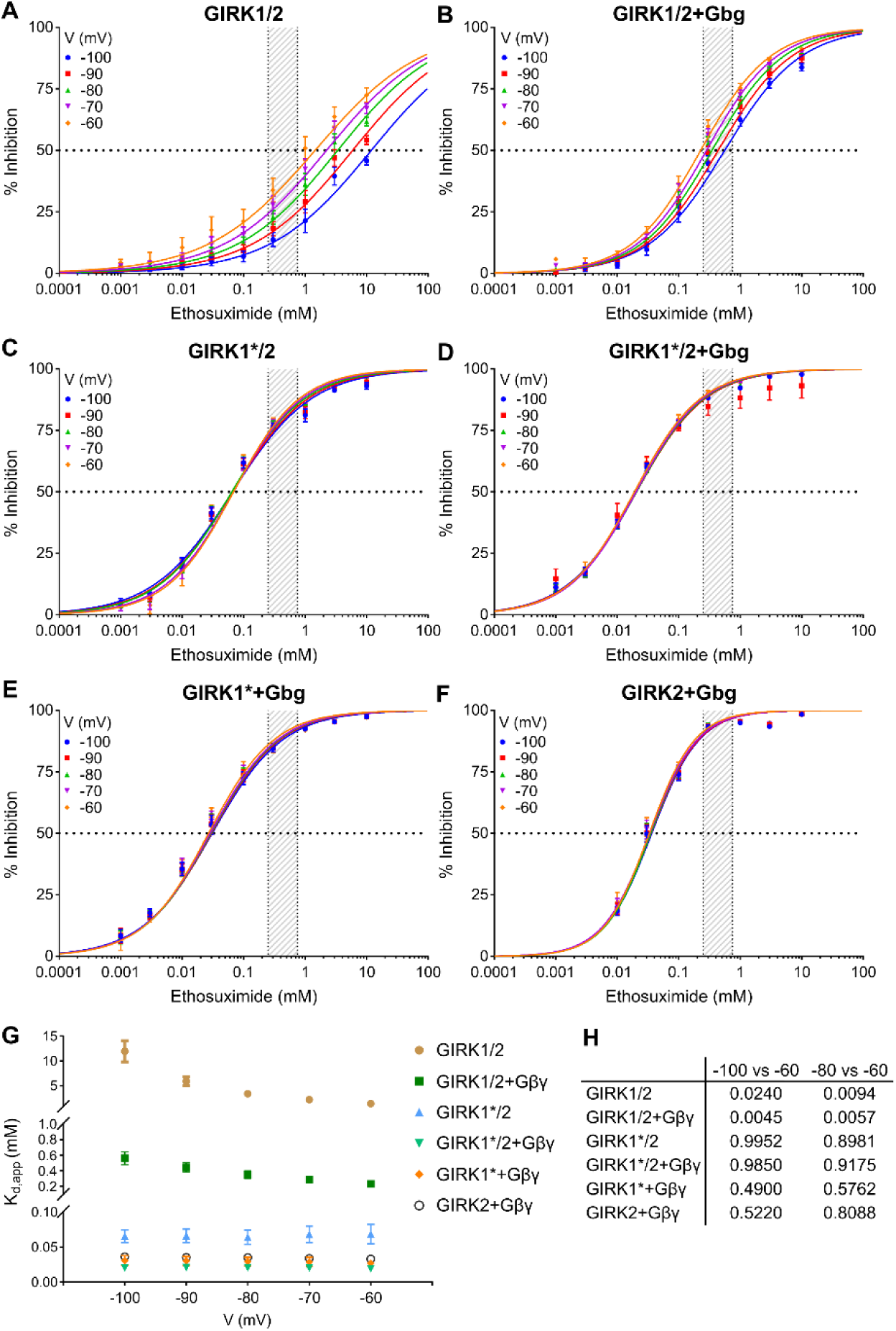
Voltage dependence of ETX block in GIRK1-containing channels. In these experiments we used mGIRK2. Currents at different voltages were determined from the I-V curves obtained by voltage ramps. **A,** GIRK1/2 channels show different apparent affinity towards ETX at different voltages. **B,** ETX block of GIRK1/2 remained sensitive to voltage with coexpressed Gβγ. **C and D,** GIRK1*/2 channels did not exhibit voltage dependence of ETX block, with or without Gβγ. **E,** GIRK1* coexpressed with Gβγ did not exhibit voltage dependence. **F,** GIRK2 coexpressed with Gβγ did not exhibit voltage dependence. **G,** comparison of K_d,app_ from all GIRK combinations shown in a-f shows that GIRK1/2 and GIRK1/2+Gβγ are blocked differently at each voltage (-100 to -60 mV, in 10 mV increments), while GIRK1*/2, GIRK1*/2+Gβγ, GIRK1*+Gβγ, and GIRK2+Gβγ had similar K_d,app_ at all voltages. Statistical analysis: two-way ANOVA (Mixed-effects model (REML)) followed by Tukey’s multiple comparisons test. **H,** statistical significance (p values) for the comparison of K_d,app_ at -100 and -80 mV vs -50 mV (same data as in g) from two-way ANOVA test.

## Supplementary Tables

**Table S1, related to Fig. 3.** Kinetic modeling of ETX block of GIRK2. The corresponding kinetic schemes (#1-5) are shown in Supplementary Fig. 3C. In all cases, n is the number of occupied Gβγ sites. In all schemes, the forward (α) and backward (β) rates for the C_4_ ↔ O transition were calculated from P_o,max_, t_o,mean_; α=111.1 s^-1^; β=1000 s^-1^. The other forward and backward rate constants were adjusted to yield EC_50_ for ETX comparable to the experimentally observed K_d,app_. Asterisk (*) indicates ETX-bound channel. **A**, summary of possible schemes of GIRK2-ETX interaction. **B**, kinetic constants used for the open-channel blocker scheme (#2). **C**, kinetic constants used for the allosteric closed channel blocker scheme (#3). **D**, kinetic constants used for the state –dependent closed channel blocker scheme (ETX was modeled to interact only with C_4_) (#4). **E**, kinetic constants used for the equal affinity closed channel blocker scheme (#5).

| Model | Assumptions | Left shift of ETX dose-response by Gβγ |
| --- | --- | --- |
| ② Open channel block | 1. ETX binds to open state only; binding of Gβγ or ETX to the channel do not affect each other's affinity. | Left shift observed |
| ③ Closed block, with allosteric coupling | 1. ETX binds to all closed states. Affinity to Gβγ increases upon ETX binding, and affinity of ETX increases upon Gβγ binding | Left shift observed |
| ④ Closed block at C4, no allosteric coupling | 1. ETX binds to only C<sub>4</sub>; binding of Gβγ or ETX to the channel do not affect each other's affinity. | Left shift observed |
| ⑤ Closed channel block, no allosteric coupling | 2. ETX binds equally to all closed states; binding of Gβγ or ETX to the channel do not affect each other's affinity. | No shift |

B
| Reaction | forward rate constant (M<sup>-1</sup>s<sup>-1</sup>) | backward rate constant (s<sup>-1</sup>) | K<sub>D</sub> (M) |
| --- | --- | --- | --- |
| C<sub>n-1</sub> + Gβγ ⇌ C<sub>n</sub> | (5-n)·1e5 (k<sub>on</sub>) | n·1 (k<sub>off</sub>) | 1e-5 (K<sub>D,Gβγ</sub>) |
| O+ETX ⇌ B | 1e5 | 1 | 1e-5 (K<sub>D,E</sub>) |

C
| Reaction | forward rate constant | backward rate constant | K<sub>D</sub> (M) |
| --- | --- | --- | --- |
| C<sub>n-1</sub> + Gβγ ⇌ C<sub>n</sub> | (5-n)·1e5 M<sup>-1</sup>s<sup>-1</sup> | n·1 s<sup>-1</sup> | 1e-5 |
| C<sub>n-1</sub>* + Gβγ ⇌ C<sub>n</sub>* | (5-n)·1e5 M<sup>-1</sup>s<sup>-1</sup> | n·0.1 s<sup>-1</sup> | 1e-6 |
| C<sub>n</sub>+ETX ⇌ C<sub>n</sub>* | 1e4 M<sup>-1</sup>s<sup>-1</sup><br>1e4 M<sup>-1</sup>s<sup>-1</sup> (k<sub>f</sub>)<br>1e4 M<sup>-1</sup>s<sup>-1</sup> (k<sub>f</sub>)<br>1e4 M<sup>-1</sup>s<sup>-1</sup> (k<sub>f</sub>)<br>1e4 M<sup>-1</sup>s<sup>-1</sup> | 1e4 s<sup>-1</sup> (k<sub>b</sub>)<br>1e3 s<sup>-1</sup> (k<sub>b</sub>)<br>1e2 s<sup>-1</sup> (k<sub>b</sub>)<br>1e1 s<sup>-1</sup> (k<sub>b</sub>)<br>1 s<sup>-1</sup> (k<sub>b</sub>) | $K_{D,E}^0 = 2e-3$<br>$K_{D,E}^4 = 1e-4$ |

D
| Reaction | forward rate constant (M<sup>-1</sup>s<sup>-1</sup>) | backward rate constant (s<sup>-1</sup>) | K<sub>D</sub> (M) |
| --- | --- | --- | --- |
| C<sub>n-1</sub> + Gβγ ⇌ C<sub>n</sub> | (5-n)·1e5 (k<sub>on</sub>) | n·1 (k<sub>off</sub>) | 1e-5 (K<sub>D,Gβγ</sub>) |
| C<sub>4</sub>+ETX ⇌ B | 1e4 | 1 | 1e-4 (K<sub>D,E</sub>) |

E
| Reaction | forward rate constant (M<sup>-1</sup>s<sup>-1</sup>) | backward rate constant (s<sup>-1</sup>) | K<sub>D</sub> (M) |
| --- | --- | --- | --- |
| C<sub>n-1</sub> + Gβγ ⇌ C<sub>n</sub> | (5-n)·1e5 | n·1 | 1e-5 |
| C<sub>n-1</sub>* + Gβγ ⇌ C<sub>n</sub>* | (5-n)·1e5 M<sup>-1</sup>s<sup>-1</sup> | n·1 s<sup>-1</sup> | 1e-5 |
| C<sub>n</sub>+ETX ⇌ C<sub>n</sub>* | 1e3 M<sup>-1</sup>s<sup>-1</sup> | 1 s<sup>-1</sup> | 1e-3 |

**Table S2.** RNA concentrations injected per oocyte, current recorded at -80 mV, and fold of activation by Gβγ (Rβγ) of channel combinations used in the experiments shown in Fig. 5.

| | RNA<br>(ng/oocyte) | I (nA)<br>at -80 mV | Rβ $\gamma$ | n | N |
| --- | --- | --- | --- | --- | --- |
| GIRK1/GIRK2<sub>WT</sub> | 0.05/0.05 | 1343±175 | | 5 | 1 |
| | 0.1/0.1 | 4225±896 | | 7 | 3 |
| GIRK1/GIRK2<sub>WT</sub>+Gβ $\gamma$ | 0.05/0.05 | 3412±358 | 2.5±0.3 | 5 | 1 |
| | 0.1/0.1 | 11123±2343 | 2.7±0.49 | 9 | 3 |
| GIRK1/GIRK3<sub>G2TM</sub> | 0.5/0.5 | 157 |  | 2 | 1 |
|  | 3/3 | 211 |  | 2 | 1 |
| | 5/5 | 130±12 | | 6 | 1 |
| GIRK1/GIRK3<sub>G2TM</sub>+Gβ $\gamma$ | 0.5/0.5 | 158 | | 2 | 1 |
| | 3/3 | 397±17 | | 8 | 1 |
| | 5/5 | 330±42 | 2.5±0.3 | 5 | 1 |
| GIRK1/GIRK3<sub>WT(Y350A)</sub> | 5/5 | 15486±287 | | 6 | 1 |
| GIRK1/GIRK3<sub>WT(Y350A)</sub>+Gβ $\gamma$ | 1/1 | 2277±455 | | 4 | 1 |
| GIRK1/GIRK2<sub>G3TM</sub> | 0.5/0.5 | 533±64 | | 9 | 3 |
| | 2/2 | 1106±188 | | 6 | 1 |
| GIRK1/GIRK2<sub>G3TM</sub>+Gβ $\gamma$ | 0.5/0.5 | 3403±641 | 6.4±1.3 | 9 | 3 |
| | 2/2 | 3944±354 | 3.6±0.3 | 3 | 1 |

**Table S3.**
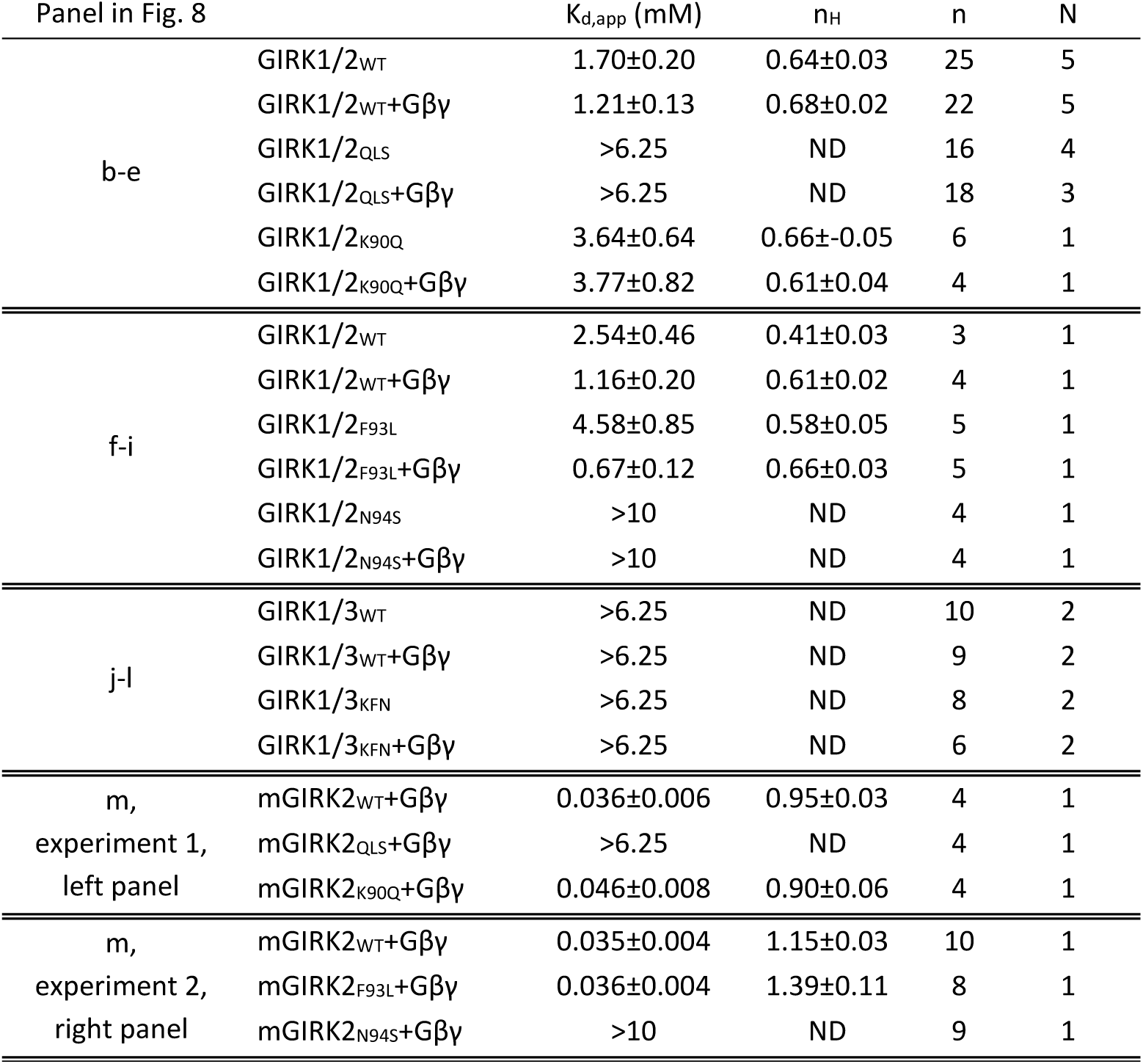
Hill equation fit parameters for the data presented in Fig. 8.

**Table S4.**
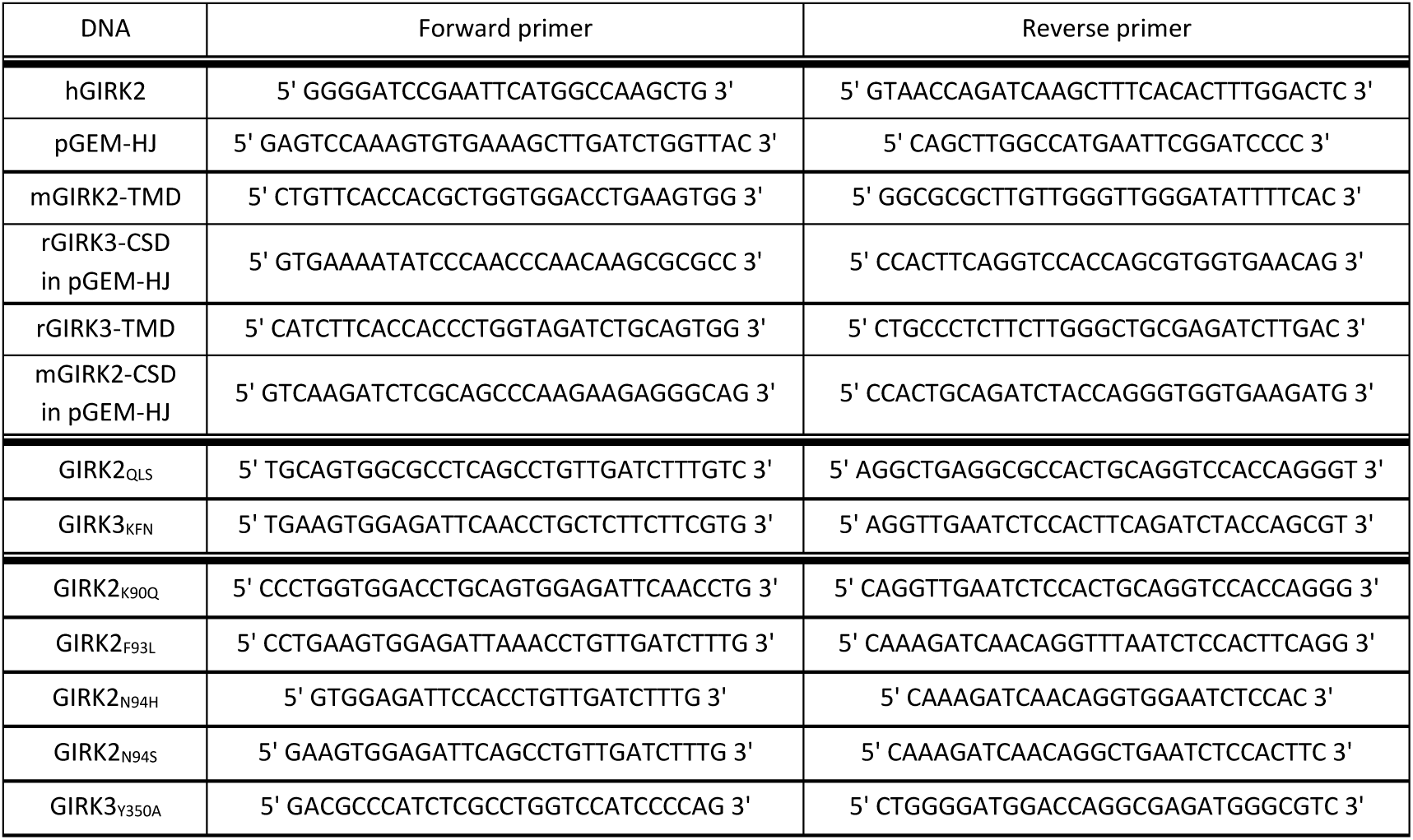
List of primers designed for subcloning of hGIRK2 into pGEM-HJ, generation of chimeras GIRK2_G3TM_ and GIRK3_G2TM_, triple mutants mGIRK2_QLS_, rGIRK3_KFN_, and point mutants mGIRK2_K90Q_, mGIRK2_F93L_, mGIRK2_N94H_, mGIRK2_N94S_, and rGIRK3_Y350A_.

**Table S5.** List of DNAs that were used for expression in Xenopus laevis oocytes.

| Gene name | Protein name | Accession number | Species |
| --- | --- | --- | --- |
| *KCNJ3* | GIRK1 | CAA72936.1 | *Rattus norvegicus* |
|  | hGIRK1<sub>F137S</sub> | P48549 | *Homo sapiens* |
| *KCNJ6* | mGIRK2 | NP_001020755.1 | *Mus musculus* |
|  | hGIRK2 | NP_002231.1 | *Homo sapiens* |
| *KCNJ9* | GIRK3 | NP_446286.1 | *Rattus norvegicus* |
| *KCNJ5* | GIRK4 | NP_058993.1 | *Rattus norvegicus* |
| *GNB1* | Gβ<sub>1</sub> | NP_786971.2 | *Bos taurus* |
| *GNG2* | Gγ<sub>2</sub> | NP_776497.1 | *Bos taurus* |
| *CACNA1C* | α1C | CAA33546.1 | *Oryctolagus cuniculus* |
| *CACNB2* | β2b | CAA45575.1 | *Oryctolagus cuniculus* |
| *CACNA2D1* | α2δ | AAA81562.1 | *Oryctolagus cuniculus* |
| *CACNA1H* | α1H | NP_001005407.1 | *Homo sapiens* |
| *CACHD1* |  | NP_001178687.1 | *Rattus norvegicus* |

